# Genome Assembly of the Roundjaw Bonefish (*Albula glossodonta*), a Vulnerable Circumtropical Sportfish

**DOI:** 10.1101/2021.09.10.458299

**Authors:** Brandon D. Pickett, Sheena Talma, Jessica R. Glass, Daniel Ence, Paul D. Cowley, Perry G. Ridge, John S. K. Kauwe

## Abstract

**Background:** Bonefishes are cryptic species indiscriminately targeted by subsistence and recreational fisheries worldwide. The roundjaw bonefish, *Albula glossodonta* is the most widespread bonefish species in the Indo-Pacific and is listed as vulnerable to extinction by the IUCN’s Red List due to anthropogenic activities. Whole-genome datasets allow for improved population and species delimitation, which – prior to this study – were lacking for *Albula* species.

**Results:** We generated a high-quality genome assembly of an *A. glossodonta* individual from Hawai‘i, USA. The assembled contigs had an NG50 of 4.75 Mbp and a maximum length of 28.2 Mbp. Scaffolding yielded an NG50 of 14.49 Mbp, with the longest scaffold reaching 42.29 Mbp. Half the genome was contained in 20 scaffolds. The genome was annotated with 28.3 K protein-coding genes. We then analyzed 66 *A. glossodonta* individuals and 38,355 SNP loci to evaluate population genetic connectivity between six atolls in Seychelles and Mauritius in the Western Indian Ocean. We observed genetic homogeneity between atolls in Seychelles and evidence of reduced gene flow between Seychelles and Mauritius. The South Equatorial Current could be one mechanism limiting gene flow of *A. glossodonta* populations between Seychelles and Mauritius.

**Conclusions:** Quantifying the spatial population structure of widespread fishery species such as bonefishes is necessary for effective transboundary management and conservation. This population genomic dataset mapped to a high-quality genome assembly allowed us to discern shallow population structure in a widespread species in the Western Indian Ocean. The genome assembly will be useful for addressing the taxonomic uncertainties of bonefishes globally.

## INTRODUCTION

Bonefishes (*Albula* spp.) are popular and economically important sportfishes found in the tropics around the globe. In the Florida Keys (Florida, USA) alone, $465 million of the annual economy is attributed to sportfishing tourism for bonefish and other fishery species inhabiting coastal flats [1]. Considering only bonefish, the sportfishing industry generates $169 million annually in the Bahamas [2, 3]. Unfortunately, population declines of bonefish have been observed around the globe, raising questions about how best to conserve bonefish and manage the associated fisheries [4]. *Albula* contains many morphological cryptic species, which, when combined with baseline data gaps, creates a significant hurdle to effective management [5-7].

All bonefish species were historically synonymized to a single species, *Albula vulpes* (Linnaeus 1758) [8], by 1940 [9-11], except for the threadfin bonefish, *A. nemoptera* (Fowler 1911) [12], which is morphologically distinct [12, 13]. Molecular testing in the last several decades has enabled specific distinctions that were not previously possible [6, 9, 14-16]. Presently, three species complexes (*A. argentea, A. nemoptera*, and *A. vulpes* complexes) contain the twelve putative albulid species, although identification remains difficult in most cases [4]. The roundjaw bonefish (Fig. 1), *A. glossodonta* (Forsskål 1775) [17], is one of seven species in the *A. vulpes* complex.

**Figure 1.**
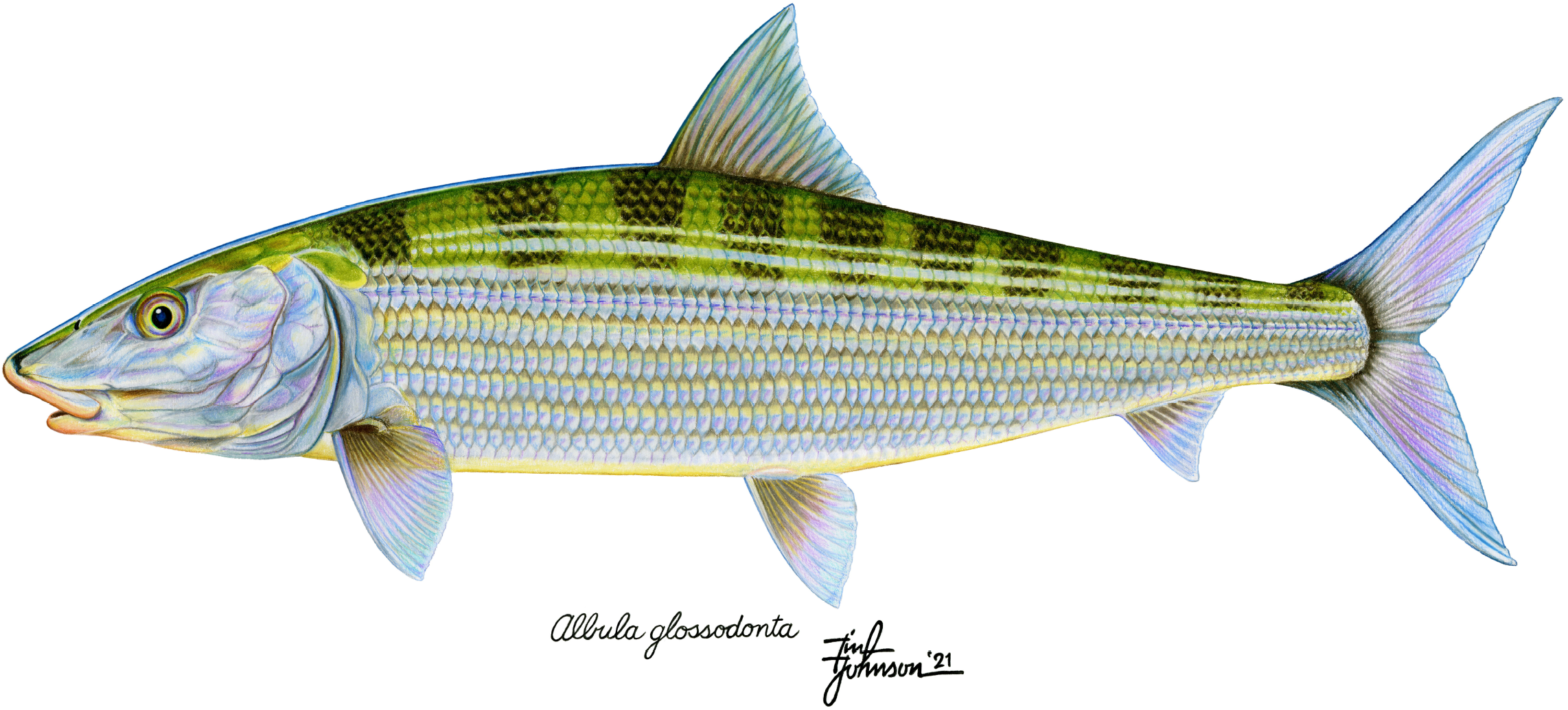
Roundjaw Bonefish (*Albula glossodonta*) adult. Quantitative morphological data for this illustration of *A. glossodonta* were obtained primarily from two articles: Hidaka et al. 2008 [163] and Shaklee and Tamaru 1981 [14]. These were then evaluated by the artist, with assistance and input from the authors, to select specific values for details such as the number of pored lateral line scales (76) and the number of rays in the pectoral (18), dorsal (16), pelvic (10), and anal fins (9). Each of these was portrayed in the illustration to be at or near the middle of the ranges reported in the aforementioned articles. While some limited information was found in the literature describing coloration and general shape, the artist found particular benefit in some excellent, detailed photographs by Derek Olthuis of samples that were both personally caught in Hawai‘i and later genetically identified as *A. glossodonta* by Dr. J. S. K. Kauwe. Illustration Copyright: Tim Johnson, used with permission.

Most of the species in the *A. vulpes* complex can be found in the Caribbean Sea and Atlantic Ocean. By contrast, *A. glossodonta* can be found throughout the Indian and Pacific Oceans; this range overlaps slightly with *A. koreana* (Kwun and Kim 2011) [18] from the *A. vulpes* complex and drastically with each species in the *A. argentea* complex [4]. *Albula glossodonta* may be distinguished genetically from other species, but morphological identification based on its more-rounded jaw and larger average size is difficult for non-experts [4, 19]. This difficulty, alongside underregulated fisheries and anthropogenic habitat loss, poses significant threats to the future of this species. In point of fact, *A. glossodonta* has been evaluated as “Vulnerable” on the International Union for the Conservation of Nature’s (IUCN) Red List of Threatened Species™ [7], and several incidents of overexploitation, including regional extirpation, have been reported [20-24].

The threat to *A. glossodonta* and other bonefish species will persist unless identification is made easier and population genomics techniques are employed to understand and identify evolutionarily significant units, areas of overlap between species, presence and extent of hybridization, and life-history traits, especially migration and spawning [4]. Genetic identification has hitherto been accomplished using only a portion of the mitochondrial cytochrome b gene and some microsatellite markers [6, 9, 15, 18, 25-32], which likely provide an insufficient taxonomic history [4, 33-35]. To contribute to a more robust capacity for identification and enable more complex genomics-based analyses, we present a high-quality genome assembly of an *A. glossodonta* individual. A transcriptome assembly was also created and was used alongside computational annotation methods to create structural and functional annotations for the genome assembly. Additionally, we present results from a population genomic analysis of *A. glossodonta* populations in Seychelles and Mauritius, two island nations that support lucrative bonefish fly fishing industries. The raw data, assembly, and annotations are available on the National Center for Biotechnology Information (NCBI) website under BioProject Accessions PRJNA668352 and PRJNA702254.

## METHODS

An overview of the methods used in this study is provided here. Where appropriate, additional details, such as the code for custom scripts and the commands used to run software, are provided in the Supplementary Bioinformatics Methods [see Additional File 1].

### Tissue Collection and Preservation

Blood, gill, heart, and liver tissues from one *A. glossodonta* individual were collected off the coast of Moloka‘i (near Kaunakakai, Hawai‘i, USA) in February 2016. Heart tissue from a second individual was also collected at the same location in September 2017. Tissue samples were flash-frozen in liquid nitrogen, and blood samples were preserved in EDTA. All samples were packaged in dry ice for transportation to Brigham Young University (BYU; Provo, Utah, USA) and stored at □80°C until sequencing. The blood sample from the first individual was used for short-read DNA sequencing. The gill, heart, and liver samples from the same individual were used for short-read RNA sequencing. The heart sample from the second individual was used for long-read sequencing and Hi□C sequencing.

For population genomic analyses, tissues (dorsal muscle samples or fin clips) were collected by fly fishing charter operators from 96 individuals of *A. glossodonta* from six coral atolls in the Southwest Indian Ocean (SWIO; Fig 1; Table S1 [Additional File 2]). All tissues were preserved in 95% EtOH at -20°C until sequencing, and thereafter cataloged and preserved in -80°C in the tissue biobank of South African Institute for Aquatic Biodiversity (Makhanda, South Africa) [36].

### Sequencing

#### DNA Sequencing

DNA was prepared for long-read sequencing with Pacific Biosciences (PacBio; Menlo Park, California, USA) [37] SMRTbell Library kits, following the protocol “Procedure & Checklist – Preparing >30 kb SMRTbell Libraries Using Megaruptor Shearing and BluePippin Size-Selection for PacBio RS II and Sequel Systems”. Continuous long-read (CLR) sequencing was performed on thirteen SMRT cells for a 10-hour movie on the PacBio Sequel at the BYU DNA Sequencing Center (DNASC) [38], a PacBio Certified Service Provider. Short-read sequencing was performed in Rapid Run mode for 250 cycles in one lane on the Illumina (San Diego, California, USA) [39] Hi-Seq 2500 at the DNASC after sonication with Covaris (Woburn, Massachusetts, USA) [40] Adaptive Focus Acoustics technology and preparation with New England Biolabs (Ipswich, Massachusetts, USA) [41] NEBNext Ultra II End Repair and Ligation kits with adapters from Integrated DNA Technologies (Coralville, Iowa, USA) [42].

#### mRNA Sequencing

RNA was prepared with Roche (Basel, Switzerland) [43] KAPA Stranded RNA-Seq kit, following manufacturer recommendations. Paired-end sequencing was performed in High Output mode for 125 cycles on the three samples together in one lane on the Illumina Hi-Seq 2500 at the DNASC.

#### Hi□C Sequencing

DNA was prepared with Phase Genomics (Seattle, Washington, USA) [44] Proximo Hi□C Kit (Animal) using the Sau3AI restriction enzyme (cut site: GATC) following recommended protocols. Paired-end sequencing was performed in Rapid Run mode for 250 cycles in one lane on the Illumina Hi-Seq 2500 at the DNASC.

#### ddRAD Library Preparation and Sequencing

We employed double digest restricted site-associated (ddRAD) sequencing to measure intraspecific genetic variation across six sampling localities in the SWIO. We extracted total DNA using Qiagen DNeasy Tissue kits per the manufacturer’s protocol (Qiagen, Inc., Valencia, California, USA) [45]. We examined the quality of DNA extractions visually using gel electrophoresis and by quantifying isolated DNA using a Qubit fluorometer (Life Technologies, Carlsbad, California, USA) [46].

We modified a protocol developed by Peterson et al. [47] to prepare samples for ddRAD sequencing. We used the rare cutter *PstI* (5′-CTGCAG-3′ recognition site) and common cutter *MspI* (5′-CCGG-3′ recognition site). We carried out double digests of 150 – 200 ng total DNA per sample using the two enzymes in the manufacturer’s supplied buffer (New England Biolabs) for 8 hours at 37°C. We randomly distributed samples from different localities across the sequencing plate to minimize bias during library preparation. We visually examined samples using gel electrophoresis to determine digestion success and then ligated barcoded Illumina adapters to DNA fragments [47]. After ligation, we pooled samples into 12 libraries and performed a clean-up using the QIAquick PCR Purification Kit. We then performed PCR using Phusion *Taq* (New England Biolabs) and Illumina indexed primers [47]. Library DNA concentration was checked using a Qubit fluorometer, followed by normalization, a second round of pooling into four libraries, and an additional QIAquick cleanup step. We then re-measured DNA concentration using a Qubit and combined equal amounts from each of the four pools into one. We analyzed this final pool using a BioAnalyzer (Agilent, Santa Clara, California, USA) [48] and performed size-selection using a Pippin Prep (Sage Science, Beverly, Massachusetts, USA) [49], selecting for fragments between 300 – 500 bp. This was followed by a final measure of concentration using a BioAnalyzer. We sent the library to the University of Oregon Genomics and Cell Characterization Core Facility [50] where concentrations were verified via qPCR before 100 bp single-end sequencing on an Illumina Hi-Seq 4000.

### Read Error Correction

#### Illumina DNA

Illumina whole-genome sequencing (WGS) reads were corrected using Quake v0.3.5 [51], which depended upon old versions of R (v3.4.0) [52] and the R package VGAM (v0.7-8) [53, 54]. Quake attempts to automatically choose a k□mer cutoff, traditionally based on k□mer counts provided by Jellyfish [55]. To generate q□mer counts instead of k□mer counts, BFCounter v0.2 [56] was used. Quake suggested a q□mer cutoff of 2.33, which was subsequently used by the correction phase of Quake. Unlike the WGS reads, the Illumina DNA reads created with the Hi□C library preparation were not corrected.

#### Illumina RNA

Illumina RNA-seq reads underwent a correction procedure using Rcorrector v1.0.2 [57]. Rcorrector automatically chooses a k□mer cutoff based on k□mer counts provided by Jellyfish [55], which Rcorrector runs automatically for the user. Alternately, Jellyfish can be run externally or bypassed with an alternate k□mer counting program, and counts can subsequently be provided to Rcorrector, which may be started at what it calls “stage 3”. We bypassed Jellyfish by using BFCounter v0.2 [56] to count k□mers. Note that Rcorrector made no changes to the reads.

#### PacBio CLRs

Several methods were attempted for the correction of the PacBio CLRs. The corrected reads from each method that did not fail were assembled, and the assembly results were used to choose the correction strategy. Ultimately, a hybrid correction strategy was employed. First, the reads were self-corrected using Canu v1.6 [58]. Second, the self-corrected reads were further corrected using Illumina short-reads (previously corrected with Quake) using CoLoRMap downloaded April 2018 [59].

### Genome Size Estimation

Genome size was estimated using a k□mer analysis on the corrected Illumina WGS reads. First, the k□mer coverage was estimated using ntCard v1.0.1 [60]. The k□mer coverage histogram was computationally processed to calculate the area under the curve and identify the peak to determine genome size according to the following equation: *a* / *p* = *s*, where *a* is the area under the curve, *p* is the number of times the k□mers occur (the x-value) at the peak, and *s* is the genome size.

### Genome Assembly, Polishing, and Scaffolding

Multiple assemblies were generated from various correction strategies. The final assembly was based on a hybrid correction strategy as described in the previous section “PacBio CLRs”. The assembly was created using Canu v1.6 [58]. The assembly underwent two rounds of polishing with the corrected Illumina WGS reads using RaCon v1.3.1 [61]. The polished contigs were scaffolded in a stepwise fashion using two types of long-range information: Hi-C and RNA-seq reads. Both scaffolding steps required read mapping to the contigs before determining how to order and orient contigs. The Hi-C data alignments were performed following the Arima Genomics (San Diego, California, USA) [62] Mapping Pipeline [63], which relied on bwa v0.7.17-r1998 [64], Picard v2.19.2 [65], and SAMtools v1.6 [66]. BEDTools v2.28.0 [67] was used to prepare the Hi-C alignments for scaffolding. The RNA-seq data were aligned using HiSat v0.1.6-beta [68]. Scaffolding was performed for the Hi-C and RNA-seq data, respectively, with SALSA, downloaded 29 May 2019 [69, 70], and Rascaf, downloaded June 2018 [71]. Assembly continuity statistics, e.g., N50 and auNG [72], were calculated with caln50 downloaded 10 April 2020 [73] and a custom Python [74] script. Assembly completeness was assessed using single-copy orthologs with BUSCO v4.0.6 [75] and OrthoDB v10 [76] (Table S2 [Additional File 2]).

### Transcriptome Assembly

The transcriptome was assembled from Illumina RNA-seq reads from all three tissues (i.e., gill, heart, and liver). The raw reads were used because Rcorrector did not modify any bases, thus making the raw reads and the “corrected” reads identical. The transcripts were assembled using Trinity v2.6.6 [77]. Assembly completeness was assessed using single-copy orthologs with BUSCO v4.0.6 [75] and OrthoDB v10 [76] (Table S2 [Additional File 2]).

### ddRAD Sequence Assembly and Filtering

We assembled all ddRAD data using the program *ipyrad* v0.9.31 [78]. The input parameters for *ipyrad* are included in the supplementary materials (Table S3 [Additional File 2]). All *A. glossodonta* reads were mapped to the genome assembly. In step one of the *ipyrad* workflow, we demultiplexed sequences by identifying individual sample barcode sequences and restriction overhangs. During step two, we trimmed barcodes and adapters from reads, which were then hard-masked using a *q*-score threshold of 20 and filtered for a maximum number of undetermined bases per read. In step three we clustered reads with a minimum depth of coverage of six to retain clusters in the ddRAD assembly. During step four, we jointly estimated sequencing error rate and heterozygosity from site patterns across the clustered reads assuming a maximum of two consensus alleles per individual. In step five, we determined consensus base calls for each allele using the parameters from step four and filtered for a maximum number of undetermined sites per locus. During step six, we clustered consensus sequences and aligned reads for each sample. During step seven, we filtered the data by the maximum number of alleles per locus, the maximum number of shared heterozygous sites per locus, and other criteria [78] and formatted output files for downstream analyses. We included all loci shared by at least 10 individuals.

We performed additional filtering steps after running *ipyrad* to account for missing data and rare alleles. Using VCFtools v0.1.16 [79] and BCFtools v1.6 [66], we removed individuals missing more than 98% of genotype calls. We retained only biallelic single nucleotide polymorphisms (SNPs) and removed (i) indels, (ii) loci with minor allele frequencies < 0.05 to exclude singletons and false polymorphic loci due to potential sequencing errors, (iii) alleles with a minimum count < 2, and (iv) loci with high mean depth values (> 100). We then implemented an iterative series of filtering steps based on missing data and genotype call rates to maximize genomic coverage per individual (Table S4 [Additional File 2]) [80]. Thereafter, we removed loci out of Hardy-Weinberg Equilibrium to filter for excess heterozygosity. We then used PLINK v1.9 [81] to perform linkage disequilibrium pruning by calculating the squared coefficient of correlation (*r*^2^) on all SNPs within a 1 kb window [82]. We removed all SNPs with an *r*^2^ value greater than 0.6.

### Computational Annotation of Assembled Genome

The MAKER v3.01.02-beta [83] pipeline was used to annotate the assembly. With minor modifications (see Supplementary Bioinformatics Methods, Additional File 1), annotation proceeded according to the process described in the most recent Maker Wiki tutorial [84]. A custom repeat library was created using RepeatModeler v1.0.11 [85]. The transcriptome assembly, genome assembly, and proteins from UniProtKB Swiss-prot [86, 87] were used as input to MAKER to create initial annotations. Gene models based on these annotations were used to train the following *ab initio* gene predictors: AUGUSTUS v3.3.2 [88, 89] and SNAP downloaded 3 June 2019 [90]. AUGUSTUS was trained using BUSCO [75] as a wrapper; SNAP was trained without a wrapper. Genemark-ES v4.38 [91-93] was also trained on the assembled genome. These models were all provided to MAKER for a second round of structural annotation. The gene models based on those annotations were filtered with gFACs v1.1.1 [94] and again provided to AUGUSTUS and SNAP. As Genemark-ES does not accept initial gene models, it did not need to be run again. The gene models from the *ab initio* gene predictors were again provided to MAKER for a third and final round of annotation. Functional annotations were added using MAKER accessory scripts, the BLAST+ Suite v2.9.0 [95, 96], and InterProScan v5.45-80.0 [97, 98]. The annotations in GFF3 format were validated with GenomeTools v1.6.1 [99] and manually curated to adhere to GenBank submission guidelines.

### Statistical Analysis of Population Genomic Data

#### Detection of Loci under Selection

Before conducting population genomic analyses, we performed outlier tests to identify loci putatively under selection, which are generally identified by a significant difference in allele frequencies between populations [100]. Specifically, we implemented two outlier detection methods that accommodate missing data: *pcadapt* v4.1.0 [100] and BayeScan v2.1 [101]. The assumption behind *pcadapt* is that loci associated with population structure, ascertained via principal component analysis (PCA), are under selection [100]. *pcadapt* is advantageously fast and able to handle large numbers of loci. The number of principal components (*K*) was chosen based on visualization of a scree plot of the eigenvalues of a covariance matrix. Once the *K*-value was chosen, the Mahalanobis distance (*D* test statistic) was calculated using multiple linear regression of the number of SNPs versus *K* [100, 102]. To account for false discovery rates, the *p*-values generated using the Mahalanobis distance *D* were transformed to q-values using the R v3.6.3 [52] *q-value* package v2.15.0 [103] with the cut-off point (α) set to 10% (0.1).

BayeScan measures allele frequencies between different populations and identifies loci that are perceived to be undergoing natural selection based on their *F*_ST_ values [104, 105]. The method applies linear regression to generate population- and locus-specific *F*_ST_ estimates and calculates subpopulation *F*_ST_ coefficients by taking the difference in allele frequency between each population and the common gene pool. BayeScan incorporates uncertainties in allele frequencies due to small sample sizes, as well as imbalances in the number of samples between populations [101]. We assigned each of the six sampling localities as a population. Our analyses were based on 1:50 prior odds and included 100,000 iterations and a false discovery rate of 10%. We used the default values for the remaining parameters and visualized results in R v3.6.3 following the developer’s manual [106]. After running both *pcadapt* and BayeScan, we used R to assess the number of outliers identified by both programs and subsequently removed outlier loci to generate a neutral dataset for downstream analyses.

#### Population Structure and Genetic Differentiation

To examine population structure, we used a model-based clustering method to reconstruct the genetic ancestry of individuals using sparse nonnegative matrix factorization (sNMF) and least-squares optimization. Model-based analyses were performed using the package *LEA* v2.6.0 [107] in R. The sNMF function in *LEA* estimates the number of ancestral populations and the probability of the number of gene pools from which each individual originated by calculating an ancestry coefficient and investigating the model’s fit through cross-entropy criterion [108]. We calculated and visualized cross-entropy scores of *K* population clusters ranging from 1–10 with 10 replicates. To complement sNMF, we also used principal component analysis (PCA), a distance-based approach based on variation in allele distributions, implemented in VCFtools v0.1.16 [79]. For sNMF and PCA analyses, no populations were assigned *a priori*. We assigned each of the six sampling localities as populations for subsequent visualization, grouped into four “island groups” based on the proximity of some of the atolls that comprised our sampling localities (Fig. 2). The five Seychelles atolls we sampled were spread amongst three separate clusters of islands that are commonly referred to as the “outer island groups” due to the geographic locations of these outlying coralline islands relative to the densely-populated, granitic “inner islands” of the Seychelles Archipelago. The island groups consisted of (i) Amirantes (St. Joseph’s Atoll), (ii) Farquhar (Farquhar and Providence Atolls), (iii) Aldabra (Aldabra and Cosmoledo Atolls), as well as (iv) Mauritius (St. Brandon’s Atoll; Table S1 [Additional File 2]). We computed summary statistics in R v3.6.3, including pairwise *F*_ST_ estimates (StAMPP v1.6.1 [109]), isolation by distance via the Mantel Rand test (adegenet v2.1.3 [110]), and expected and observed heterozygosity (hierfstat v0.5-7 [111]) to compare genetic diversity and differentiation between the four island groups.

**Figure 2.**
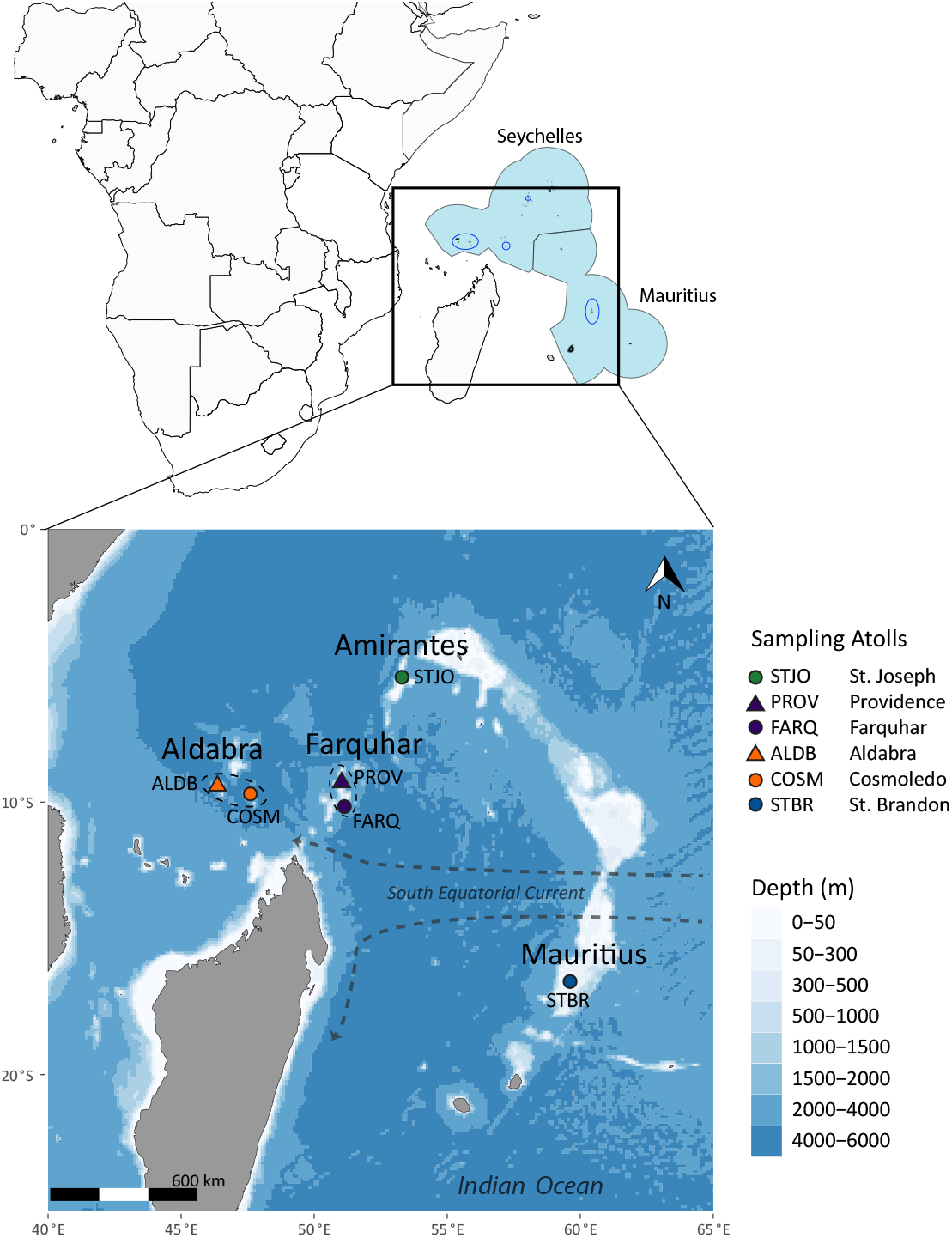
Sampling localities for *A. glossodonta* population genomic analysis. The upper panel shows the marine boundaries for the Seychelles and Mauritius in light blue. Locations of sampling sites are indicated by dark blue ovals. The lower panel shows the atolls comprising the four island groups: Amirantes, Farquhar, Aldabra, and Mauritius.

## RESULTS

### Sequencing

#### DNA Sequencing

Paired-end, short-read sequencing (Illumina) yielded 109.5M pairs of reads comprised of 53.86Gbp. The mean and N50 read lengths were 245.981 and 250, respectively. Continuous long-read sequencing (PacBio) generated 9.5M reads with a total of 69.85Gbp. The mean and N50 read lengths were 7,352.726 and 13,831, respectively. The longest read was 103,889bp. The read length distribution is plotted in Figure 2. Result summaries for both sequencing runs are available in Table 1.

**Table 1.**
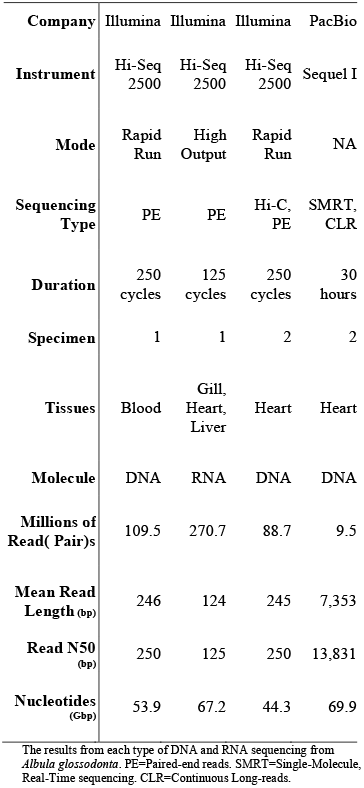
Sequencing Information.

#### mRNA Sequencing

RNA-seq from the three tissues (i.e., gill, heart, and liver) generated 270.7M pairs of reads totaling 67.2Gbp. The gill tissue yielded 107.7M pairs of reads, with a total of 26.7Gbp. The heart tissue generated 19.6Gbp across 78.8M pairs of reads. The 84.2M pairs of reads from the liver tissue were comprised of 20.9Gbp. Across all three tissues, the mean and N50 read lengths were 124.122 and 125, respectively. The combined results from all three tissues are summarized in Table 1.

#### Hi□C Sequencing

Sequencing yielded 88.7M pairs of reads comprised of 44.28Gbp. The mean and N50 read lengths were 249.493 and 250, respectively. A summary of these results is presented in Table 1.

#### ddRAD sequencing

After data processing using *ipyrad*, we recovered a mean of 114,324 reads per individual for *A. glossodonta* and an average of 107,105 loci per individual. Following filtering for missing data, minor allele frequency, and linkage disequilibrium, the dataset contained 66 individuals and 38,355 SNPs. BayeScan, being a more conservative outlier detection method than *pcadapt*, did not identify any outliers; we thus used only outlier detection results from *pcadapt*. Subsequent removal of *pcadapt* outliers (N = 155) resulted in a neutral dataset containing 38,200 SNPs with 9% missing data.

### Read Error Correction

#### Illumina DNA

When Quake corrects paired-end reads, three outcomes are possible for each pair of reads: (i) both reads are either already correct or correctable, (ii) one read is either correct or correctable and the other is low-quality, or (iii) both reads are low-quality. Of the original 218.96M reads (109.5M pairs of reads), Quake corrected 62.7M of them and removed 51.6M of them. 5.97M pairs of reads were discarded because both reads were rated as erroneous. 39.6M pairs of reads had one read that was correct or correctable and one read that was low-quality; these were also discarded. The remaining 63.88M pairs of reads were either correct or correctable and were kept in the final read set containing 29.11Gbp of sequence.

#### Illumina RNA

No corrections were made to the RNA-seq reads by the error correction software.

#### PacBio CLRs

The dual-correction strategy (self-correction followed by hybrid-correction) reduced the number of reads from 9.5M to 2.79M and the total number of bases from 69.85Gbp to 36.79Gbp. The mean and N50 read lengths were changed from 7,354 and 13,831 to 13,193 and 15,483, respectively. The longest read was 63,271 bases. The distribution of read lengths can be viewed in Fig. 3.

**Figure 3.**
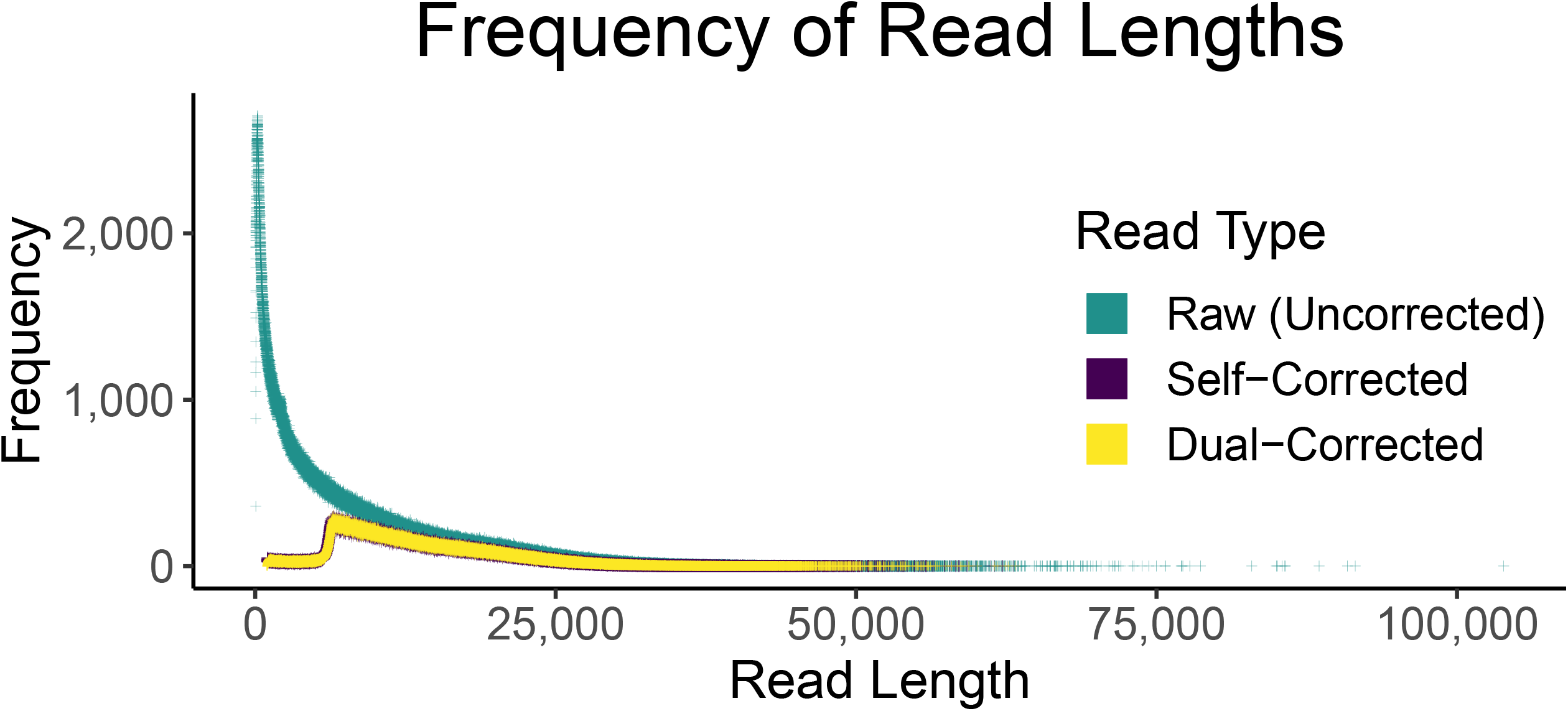
Frequency of Pacific Biosciences Read Lengths. The change in read length distribution is demonstrated as reads are corrected. The dramatic shift from raw to corrected reads is evident. The self-corrected (purple) data points are slightly larger than the dual-corrected (yellow) data points to make the purple distribution visible, the size has no meaning.

### Genome Size Estimation

The genome size was estimated to be approximately 0.933Gbp as a result of the k□mer analysis, which was consistent with the authors’ expectations based on two closely related elopomorph species [112, 113].

### Genome Assembly, Polishing, and Scaffolding

The initial assembly from Canu was comprised of 3.8K contigs with a total assembly size of 1.05Gbp. The mean contig length, N50, NG50, and maximum contig length were 276.2Kbp, 3.6Mbp, 4.7Mbp, and 28.2Mbp, respectively. The L50 was 57, and the LG50 was 43. The auNG was 8.17M. After two rounds of polishing these contigs with the corrected Illumina WGS reads using RaCon, the assembly statistics changed only marginally. The number of contigs, L50, and LG50 were unchanged. The assembly size decreased by 318.7Kbp (0.03%). The mean contig length, N50, NG50, and maximum contig length were reduced by 83.8bp (0.03%), 1.3Kbp (0.04%), 1.5Kbp (0.03%), and 3.8Kbp (0.01%), respectively. The auNG decreased by 2Kbp (0.02%).

The scaffolding with the Hi-C data joined some polished contigs together, reducing the sequence count to 3.6K (−4.69%). The number of bases, excluding unknown bases (Ns), was unchanged; however, it is important to note that when SALSA creates gaps while ordering and orienting contigs, it always uses a gap size of 500bp. The result, in this case, was adding 116Kbp of Ns, which means 232 gaps were created. These gaps were spread across 113 scaffolds. No scaffold had more than six gaps (seven contigs ordered and oriented together). The mean scaffold length, scaffold N50, scaffold NG50, and maximum scaffold length increased by 13.6Kbp (4.92%), 3.8Mbp (106.25%), 5.79Mbp (121.90%), and 14.1Mbp (49.85%), respectively. Coupled with these increases were decreases of 29 (50.88%) and 22 (51.16%) in the L50 and LG50, respectively. The auNG increased to 14.1M (+72.81%). The quality of the Hi□C scaffolding can be visualized (Fig. 4) via a contact matrix generated by PretextMap [114] and PretextView [115].

**Figure 4.**
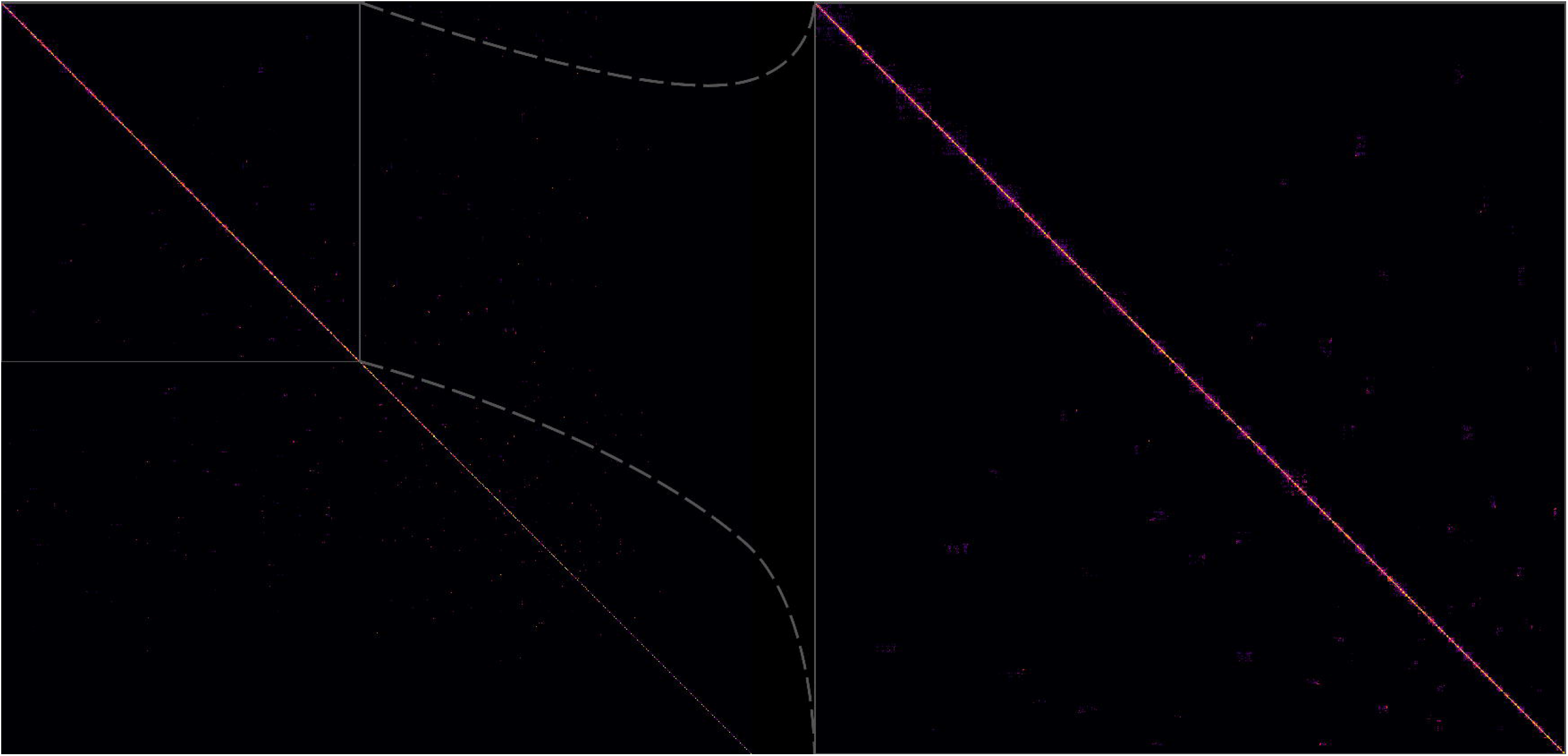
Hi-C Contact Matrix showing Scaffolding Correctness. In the context of scaffolding, Hi-C contact matrices show how correct the scaffolds are. Off-diagonal marks, especially those that are bright and large, are evidence of mis-assembly and/or incorrect scaffolding. The interpretations of the lighter and smaller off-diagonal marks in this image are ambiguous because the assembly is unphased with some relatively short contigs/scaffolds. Additional detail may be viewed by zooming in on the high-resolution image.

The genome assembly was further improved by scaffolding with RNA-seq data. As expected, the magnitude of the changes between sets of scaffolds was smaller than what was observed between contigs and scaffolds. The total number of sequences was reduced by 176 to 3.4K (−4.69%). The number of known bases was again unchanged; however, it is important to note that when Rascaf orders and orients contigs (or other scaffolds) it always inserts a gap of 17bp to represent gaps of unknown size. Rascaf added 179 new gaps (3,043 unknown bases) across 148 sequences. Three gaps (1,500 unknown bases) from SALSA were removed, but the rest remained unchanged. The most gaps added to a single sequence by Rascaf was five. The sequence with the most total gaps (from either source) had seven gaps (six from Hi-C), thus eight contigs were joined together.

This resulting set of scaffolds (which also includes all the contigs that were not joined to another contig in some way) had a mean length of 304.5Kbp (+5.11% from the Hi-C only value) and a maximum length of 42.29Mbp (+0.08%). The N50 and NG50 increased to 7.97Mbp (+7.04%) and 14.49Mbp (+37.58%), respectively. Decreases to 26 (−7.14%) and 20 (−4.76%) were observed for L50 and LG50, respectively. The auNG increased to 14.7M (+4.37%). Table 2 summarizes the assembly continuity statistics, and the area under the NG-curve (auNG) is visualized in Fig. 5.

**Table 2.**
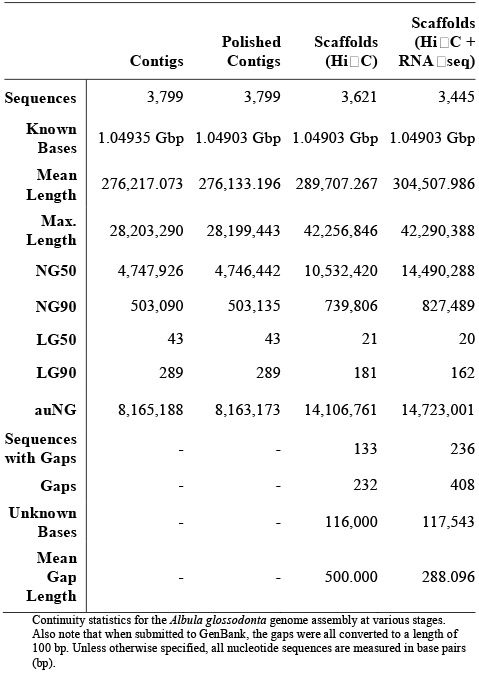
Continuity Statistics.

**Figure 5.**
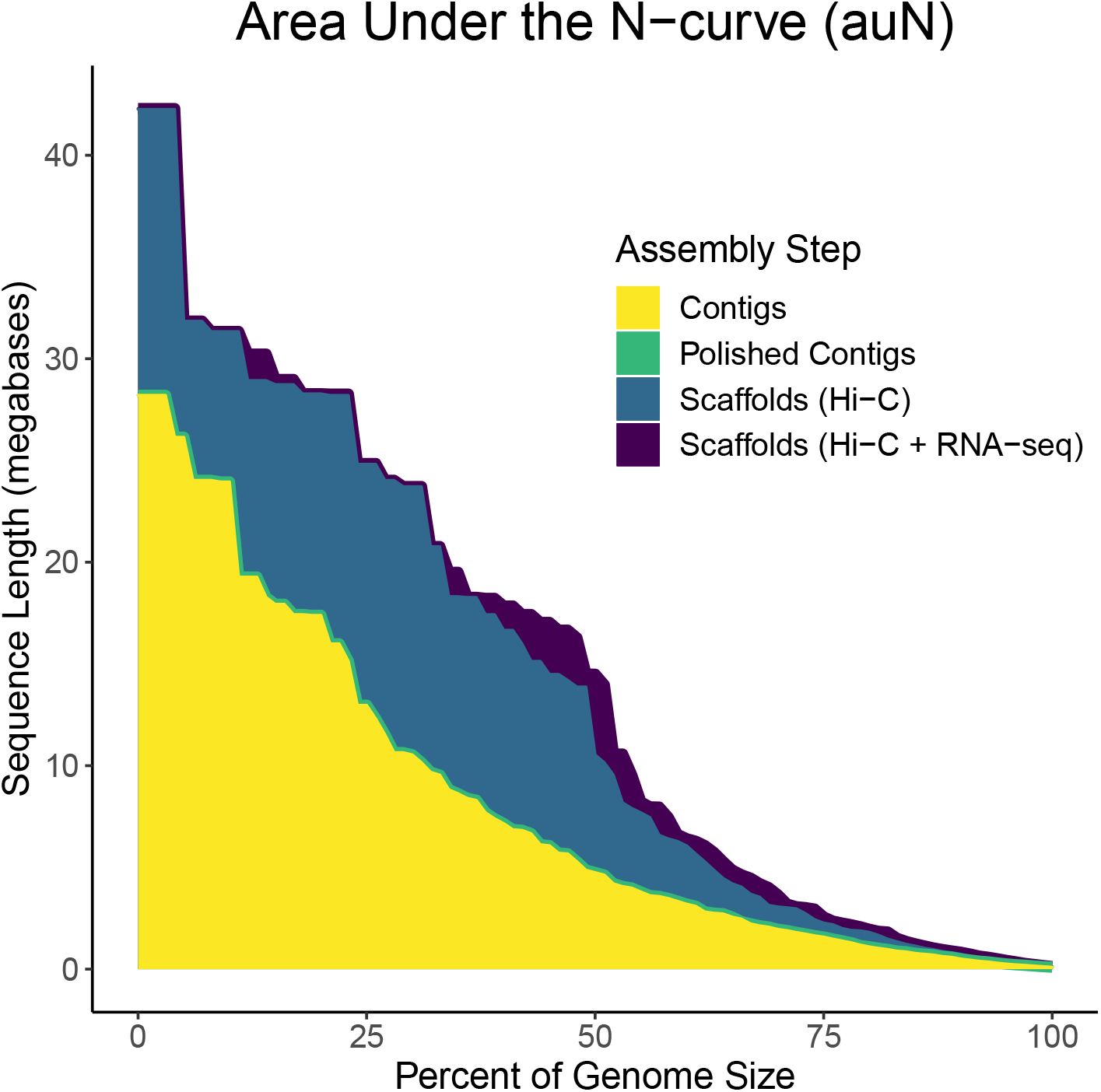
Area Under the N-curve (auNG) for each Assembly Step. The N-curve and the area under it are plotted for each major step of the assembly: contigs, polished contigs, scaffolds from only Hi-C data, and scaffolds from both Hi-C and RNA-seq data. The auNG for the polished contigs (green) is very similar to the contigs (yellow). Most of the curve was completely blocked by the contigs (yellow) curve. To show that the polished contigs (green) share nearly the same curve, the line was plotted more thickly so it can just barely be seen. Similarly, the Hi-C + RNA-seq scaffolds (purple) curve is very similar to the Hi-C only scaffolds (blue) curve. In this case, differences are more apparent. In certain places, e.g., at the highest peak, the Hi-C + RNA-seq scaffolds (purple) are plotted more thickly so it can be seen behind the Hi-C only scaffolds (blue).

The assembly completeness, as assessed with single-copy orthologs, was also evaluated at each stage (Table S2 [Additional File 2]). The results suggest that the modifications made to the primary Canu-based assembly from polishing and scaffolding did not significantly impact the correct assembly of single-copy orthologs. The final set of scaffolds had 3,481 complete single-copy orthologs (95.6% of 3,640 from the ODB10 Actinopterygii set). Of these 88.4% (3,076) were present in the assembly only once, and 11.6% (405) were present more than once. Twenty-five (0.7%) and 135 (3.7%) single-copy orthologs were fragmented in and missing from the assembly, respectively.

### Transcriptome Assembly

The transcriptome assembly generated by Trinity was comprised of 455K sequences with a mean sequence length of 1,177bp. The N50 and L50 were 2.6Kbp and 56K, respectively. The N90 and L90 were, respectively, 410bp and 270K. Of the 3,640 single-copy orthologs in the ODB10 Actinopterygii set, 86.4% (3,144) were complete; 39.5% (1,241) of which were present only once in the transcript set. 128 (3.5%) single-copy orthologs were fragmented in the transcript set, 368 (10.1%) were missing. (See Table S2 [Additional File 2])

### Computational Genome Annotation

Computational structural and functional annotation yielded 28.3K protein-coding genes. Of these, 17.2K and 15.6K have annotated 5′ and 3′ UTRs, respectively. 1.8K tRNA genes were also identified. The annotations are available with the assembly on GenBank.

### Population Genomic Analysis

Cross-entropy scores generated by the model-based population differentiation analysis, sNMF, provided support for a single population of *A. glossodonta* across all localities. However, individual ancestry plots generated by sNMF showed evidence of genetic differentiation in individuals from Mauritius (St. Brandon’s Atoll), compared to the Seychelles sites (Fig. 6A). This differentiation was corroborated by PCA visualization of the first two principal components, where St. Brandon’s Atoll individuals clustered separately from the four Seychelles island groups (Fig. 6B). Together, both population differentiation analyses indicated weak geographic population structure across all sampling localities, with reduced gene flow between St. Brandon’s Atoll and the Seychelles sites.

**Figure 6.**
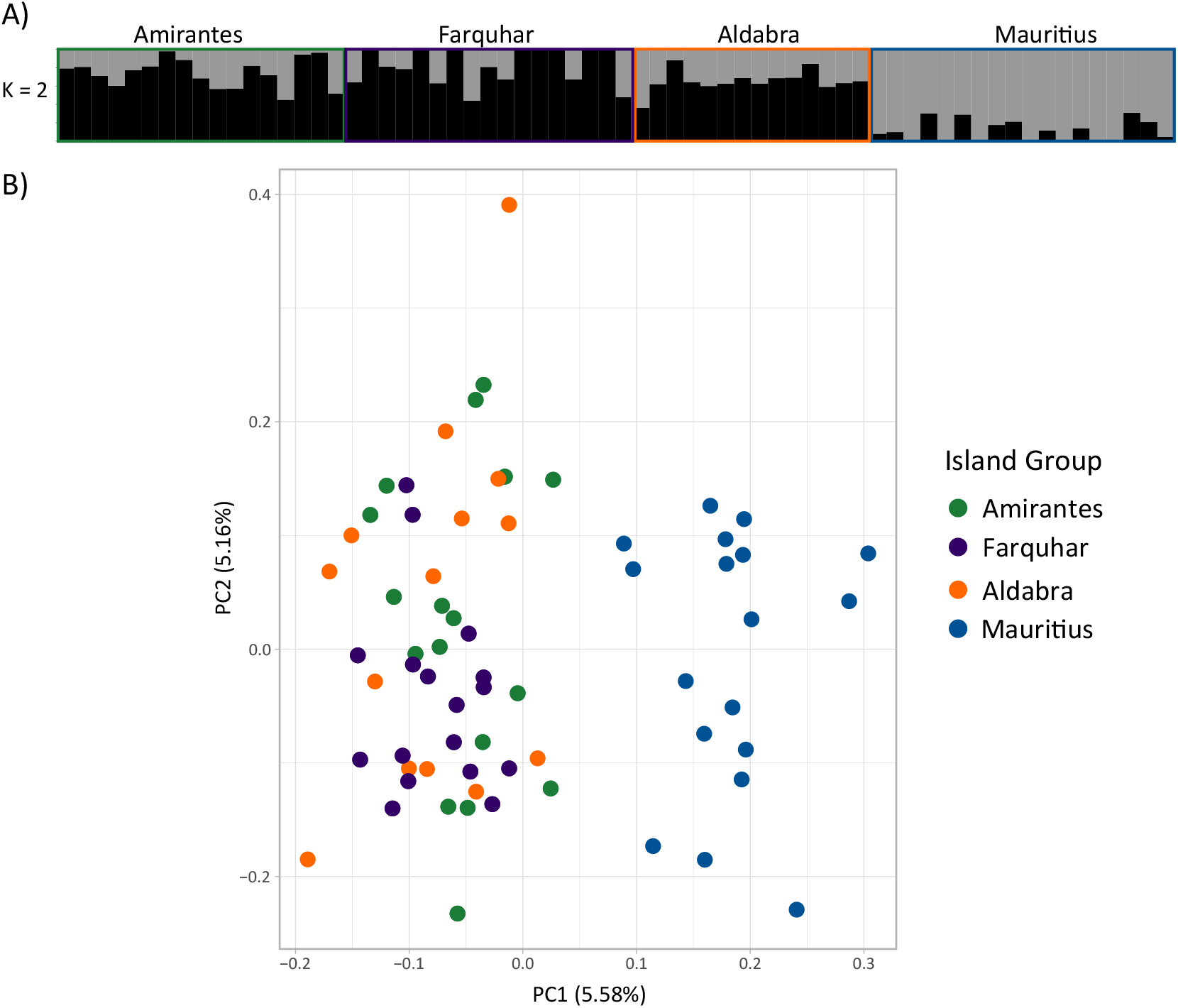
Population Differentiation Analyses. Weak geographic population structure is present across all sampling localities, with reduced gene flow between St. Brandon’s Atoll and the Seychelles sites. Island groups are colored as in Fig. 2. **(A)** Individual ancestry plots generated using sNMF, indicating *K* = 2 putative populations. **(B)** Principal component analysis biplot showing the first two principal components.

Pairwise *F*_ST_ results also indicated greater genetic differentiation between St. Brandon’s Atoll and all other island groups (Table 3). Estimates of observed and expected heterozygosity were similar across island groups (Table S5 [Additional File 2]), suggesting no differences in genetic diversity between sampling localities and providing no evidence for distinguishing metapopulation processes such as inbreeding. A test of isolation by distance between sampling sites was not significant (*p* = 0.1501).

**Table 3.**
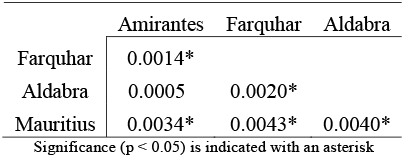
Pairwise *F*_ST_ comparisons by island group.

## DISCUSSION

*Albula glossodonta* is an important fishery species in the Indo-Pacific for both subsistence and recreational purposes [20, 30, 116, 117]. Given this species’ current “Vulnerable” IUCN status [7, 118] amidst recent taxonomic uncertainties [4], understanding patterns of gene flow and population structure in *A. glossodonta* is important for fisheries management [30, 119].

We observed a genetically homogenous population of *A. glossodonta* across five island atolls in the Seychelles Archipelago, with limited gene flow between Seychelles and Mauritius. Unlike highly migratory species such as eels (Anguillidae), which are close relatives of bonefishes, adult bonefishes are known for high site fidelity with relatively short migrations (∼10-100 km) [117, 120, 121]. We hypothesized that adult bonefishes would not migrate between the Seychelles islands, or between the Seychelles and St. Brandon’s Atoll in Mauritius, since these distances span 400–2,000 km. Consequently, the observed trend of genetic homogeneity across the Seychelles is likely not a result of adult long-distance migrations, but rather pelagic larval dispersal, the primary dispersal mechanism for bonefishes [32, 121-123]. Bonefish larvae, also referred to as leptocephali, have a long pelagic larval duration ranging from 41–72 days, which enables them to drift long distances with ocean currents [21, 124]. The estimated pelagic larval duration for *A. glossodonta* is 57 days, based on observations of individuals from French Polynesia in the South Pacific [21]. The Seychelles islands are located in the South Equatorial Current, which flows westwards from the Indian Ocean towards the eastern coast of continental Africa, enabling larvae to be transported across the Seychelles islands, even across depths exceeding 4000 m (Fig. 2) [125, 126].

Genetic homogeneity is not always an outcome of long pelagic larval duration, as demonstrated by *Anguilla marmorata*, for which 2–5 stocks were identified in the Indo-Pacific [127, 128], and *A. glossodonta*, where putative stocks between the Indian and Pacific Oceans were suggested [119]. Indeed, we found evidence of restricted gene flow between the Seychelles sampling sites and St. Brandon’s Atoll, Mauritius, which is ∼1500–2000 km from the Seychelles Islands (Fig. 2). This genetic structuring was unexpected, given the long pelagic larval duration of *A. glossodonta*. However, there is evidence of limited gene flow between Seychelles and Mauritius in other marine fish species with pelagic larvae, such as *Lutjanid kasmira* [129], *Lethrinus nebulosus* [130], and *Pristipomoides filamentous* [131].

We attribute the observed genetic structure between Seychelles and St. Brandon’s Atoll to the ocean currents in the southwestern Indian Ocean and their role in larval transport [132, 133]. St. Brandon’s Atoll is in the direct path of one of the bifurcated arms of the South Equatorial Current as it passes through the Mascarene Plateau [125, 134]. The South Equatorial Current pushes water westward, which may create a barrier to gene flow to islands south of Seychelles such as Mauritius and Réunion [130, 131, 134]. Although there are currently no bonefish – or even elopomorph – larval dispersal models for the Indian Ocean, pelagic larval dispersal simulation models of coral species in the southwestern Indian Ocean corroborate the biogeographic break between Seychelles and Mauritius, suggesting connectivity is limited even when the pelagic larval duration is between 50–60 days [125, 134]. However, these models considered coral larvae, which are completely reliant on currents for their dispersal [122, 134, 135]. Whilst the dispersal behavior of *A. glossodonta* larvae is unknown, we speculate that, similar to eels (Anguillidae; which also have long pelagic larval durations), bonefishes could disperse greater distances than passive corals by having the ability to swim (e.g., *Anguilla japonica* [136]) or may even take part in vertical migrations (e.g., *Anguilla japonica* [137, 138]). While officially undescribed, swimming ability in bonefish leptocephali has been observed [139], and vertical migrations have previously been theorized [122, 140].

Genome-wide datasets have enabled researchers to better-delineate population connectivity across seascapes for marine species where conventional markers (e.g., mtDNA, microsatellites) have not provided sufficient genomic resolution [127, 141, 142]. Such advances in genomic sequencing have altered our view of population connectivity in other marine fishes such as yellowfin tuna (*Thunnus albacores* [143]) and the American eel (*Anguilla rostrata* [144]). These studies, including ours, highlight the power of large genomic datasets for investigating connectivity in open-ocean environments containing few, if any, natural barriers that were traditionally thought to drive population structure. Although there has been a rapid increase in the number of studies using next-generation sequencing datasets for marine fishes, few studies to date have employed the use of genomic datasets on elopomorphs, and none on bonefish [144-146].

## Conclusions

This is the first genome assembly and annotation for an albulid species, as well as the first use of a genome-wide single-nucleotide polymorphism dataset to investigate population structure for *Albula glossodonta* or any bonefish species in the Indian Ocean. Individuals of *A. glossodonta* were genetically homogenous across four coralline island groups in the Seychelles Archipelago, but they showed evidence of genetic differentiation between the Seychelles and Mauritius (St. Brandon’s Atoll). These patterns of connectivity are likely facilitated by pelagic larval dispersal, which is presumed to be strongly shaped by currents in the southwestern Indian Ocean. Only with high-resolution genomic data were we able to discern this pattern of population structure between Seychelles and Mauritius. Our dataset serves as a valuable resource for future genomic studies of bonefishes to facilitate their management and conservation.

## DATA AVAILABILITY

The raw reads, genome assembly, and annotations are available under BioProject PRJNA668352 and BioSamples SAMN16516506-SAMN16516510 and SAMN17284271. The ddRAD reads are available under BioProject PRJNA702254, BioSamples SAMN18012541-SAMN18012606.

## AUTHOR CONTRIBUTIONS

**PDC:** Conceptualization; Funding Acquisition; Investigation; Supervision; Resources; Writing - Review & Editing. **DE:** Methodology; Validation; Writing - Original Draft Preparation; Writing - Review & Editing. **JRG:** Conceptualization; Formal Analysis; Investigation; Supervision; Methodology; Visualization; Writing - Original Draft Preparation; Writing - Review & Editing. **JSKK:** Conceptualization; Funding Acquisition; Investigation; Supervision; Resources; Writing - Review & Editing. **BDP:** Conceptualization; Data Curation; Formal Analysis; Investigation; Methodology; Software; Visualization; Writing - Original Draft Preparation; Writing - Review & Editing. **PGR:** Funding Acquisition; Supervision; Resources; Writing - Review & Editing. **ST:** Investigation; Resources; Writing - Original Draft Preparation; Writing - Review & Editing.

## ACKNOWLEDGEMENTS

We thank the artist, Tim Johnson [147], for creating the beautiful illustration (Fig. 1). We thank the Brigham Young University DNA Sequencing Center [38] and Office of Research Computing [148] for their continued support of our research. We thank Elizabeth M. Wallace, Clayton Ching, Josiah Ching, Derek Olthuis, Zachary Emig, Weston Gleave, and the fly fishing guides from FlyCastaway [149] and Alphonse Fishing Company [150], especially Daniel Hoenings and Matthieu Cosson, for the collection of samples in Hawai‘i and the western Indian Ocean. We are grateful to Taryn Bodill and Martinus Scheepers of the South African Institute for Aquatic Biodiversity [36] for laboratory assistance and Thomas Near of Yale University [151] for the use of laboratory space, funding, and equipment. We also thank the Seychelles Fishing Authority [152], the Island Conservation Society [153], the Islands Development Company Ltd. [154], the Seychelles Islands Foundation [155], the Ministry of Agriculture, Climate Change and Environment [156], and Shane and Hafiza Talma for their logistical support.

## ADDITIONAL FILES

**Additional File 1:** Supplementary Bioinformatics Methods (PDF).

**Additional File 2:** Supplementary Tables (PDF).

