## Additional File 1 for "Genome Assembly of the Roundjaw Bonefish (*Albula glossodonta*), a Vulnerable Circumtropical Sportfish"

**Supplementary Bioinformatics Methods**

An overview of the methods used in this study was provided in the main manuscript. Where appropriate, additional details, such as the code for custom scripts and the commands used to run software, are provided here.

**S.1 – Tissue Collection and Preservation**

Not applicable.

**S.2 – Sequencing**

Not applicable.

**S.3 – Read Error Correction**

*S.3.1 – Illumina DNA*

An estimate of the number of k-mers present in the reads is required to run BFCounter. This number is really just a simple math problem based on the number of reads, the length of the reads, and k-mer size according to this equation:

$$T=n(l-k+1)$$

Where *n* is the number of reads, *l* is the read length, and *k* is the k-mer size, and *T* is the total number of k-mers (not necessarily unique or distinct) present in the reads. Of course, this assumes a uniform read length. If the reads are paired-end, n is still the number of reads, not the number of pairs of reads. Since ntCard v1.0.1 (Hamid et al. 2017) was used to quickly get a picture for the k-mer coverage histogram, its reported value F0 was used instead of the equation as it is an estimate for *T*. ntCard was run according to the following command:

```
ntcard \
    -k 19\
    -t ${THREADS} \
    -p ${OUTPUT_FILE_BASE_NAME} \
    ${INPUT_FASTQ_FILES[@]}
```

To generate q‑mer counts BFCounter v0.2 (Melsted and Pritchard 2011) was used to count and dump the q-mers according to the following commands:

```
BFCounter count\
    -k 19\
-n ${TOTAL_NUMBER_OF_KMERS} \
-s ${RANDOM_SEED} \
-t ${THREADS} \
-o ${COUNTS_FILE_NAME} \
--quake \
--quality-scale=33 \
${INPUT_FASTQ_FILES[@]}

BFCounter dump\
    -k 19\
-i ${COUNTS_FILE_NAME} \
-o ${OUTPUT_FILE_NAME} \
--quake
```

Quake v0.3.5 (Kelley et al. 2010) was run in two stages where the first identifies a q-mer cutoff and the second corrects the reads based on that cutoff. The suggested q‑mer cutoff was 2.33, which was subsequently used by the correction phase of Quake. The two steps were executed according to the following commands:

```
cov_model.py \
${BFCOUNTER_DUMP_FILE}

correct \
    -k 19 -q 33 \
-m ${QMER_COUNTS_FILE} \
-o ${OUTPUT_FILE_NAME} \
-f ${INPUT_FASTQ_FILES[@]} \
-p ${THREADS} \
-c ${CUTOFF} \
-u --headers --log
```

Quake was developed quite some time ago, and the installation process was made difficult as dependencies were updated and function calls were broken. Multiple solutions likely exist to remedy the problem, but we found success by installing Quake with R v3.4.0 (https://www.r-project.org) with package VGAM v0.7-8 (https://CRAN.R-project.org/package=VGAM) (Yee and Wild 1996).

*S.3.2 – Illumina RNA*

Since no corrections were made by Rcorrector v1.0.2 (Song and Florea 2015) and the command is fairly straightforward, little additional detail is necessary. Recall that BFCounter was used instead of the built-in Jellyfish to generate the counts. Also note that this process was run separately for each tissue. The commands used are the following:

```
BFCounter count\
    -k 19\
-n ${TOTAL_NUMBER_OF_KMERS} \
-s ${RANDOM_SEED} \
-t ${THREADS} \
-o ${COUNTS_FILE_NAME} \
--quality-scale=33 \
${INPUT_FASTQ_FILES[@]}

BFCounter dump\
    -k 19\
-i ${COUNTS_FILE_NAME} \
-o ${DUMP_FILE_NAME} \

rcorrector \
    -k 19 \
-c ${DUMP_FILE_NAME} \
-od ${OUTPUT_DIR_NAME} \
-p ${INPUT_FASTQ_FILES[@]} \
-t ${THREADS}
```

*S.3.3 – PacBio CLRs*

First the process to correct the PacBio CLRs will be described. Next, the experiments with other correction strategies will be briefly described.

*S.3.3.1 – Dual Correction Strategy*

Typically, a “hybrid” correction strategy is defined as one in which more than one data type (i.e., PacBio CLRs and Illumina short reads) are employed. This differs from a “self” correction strategy in which only the PacBio CLRs are used to correct themselves. We employed a strategy that is “hybrid”, but that is not fully described by the word “hybrid”. We have referred to this strategy as “dual” correction. First, “self” correction is completed. Second, “hybrid” correction is done on the already self-corrected reads. The self-corrected reads were generated using Canu v1.6 (Koren et al. 2017) with the following command:

```
canu -correct \
    -s ${SETTINGS_FILE} \
-d ${OUTPUT_DIR_NAME} \
-p ${OUTPUT_PREFIX} \
-pacbio-raw \
${INPUT_PACBIO_READS[@]}
```

The relevant lines of the setting file are included here:

genomeSize=932813000

ovsMethod=sequential

gridEngine=slurm

The self-corrected reads were provided to CoLoRMap downloaded April 2018 (Haghshenas et al. 2016) as the “uncorrected” input reads. Please note that you will need to combine and interleave all Illumina short reads into a single file. All PacBio reads will also need to be in a single file, and the headers will need to be unique up to the first space, so some modification to the headers may be necessary. CoLoRMap is really a pipeline with a very basic wrapper script. In practice, it makes more sense to run each step in the wrapper script as separate jobs to avoid re-computing if a failure (e.g., too much RAM or time) occurs in a downstream step. If nothing else, a simple addition of logical checks can be added to the wrapper script to ensure subsequent steps aren’t run if the previous step failed. If run without any such modifications, the commands to run CoLoRMap are the following:

```
runCorr.sh \
    ${INPUT_SELF_CORRECTED_PACBIO_READS} \
${INPUT_ILLUMINA_READS} \
${OUTPUT_CORRECTED_PACBIO_READS_DIR} \
${OUTPUT_CORRECTED_PACBIO_READS_PREFIX} \
${THREADS}

runOEA.sh \
    ${INPUT_COLORMAP_CORRECTED_READS} \
${INPUT_ILLUMINA_READS} \
${OUTPUT_CORRECTED_PACBIO_READS_DIR} \
${OUTPUT_CORRECTED_PACBIO_READS_PREFIX} \
${THREADS}
```

Once the correction and overlap error extension assembly phases are completed, the now “dual” corrected reads are ready for assembly.

*S.3.3.2 – Correction Experiments*

We explored the effects on assembly continuity of several correction strategies before settling on the chosen strategy. Ignoring failed strategies due to software failures, three strategies were employed: (a) “self” correction (only PacBio CLRs, (b) “hybrid” correction (using only Illumina reads to correct the PacBio CLRs), and (c) “dual” correction (using Illumina reads to correct already self-corrected PacBio CLRs). These correction strategies are described visually in the following flow chart:

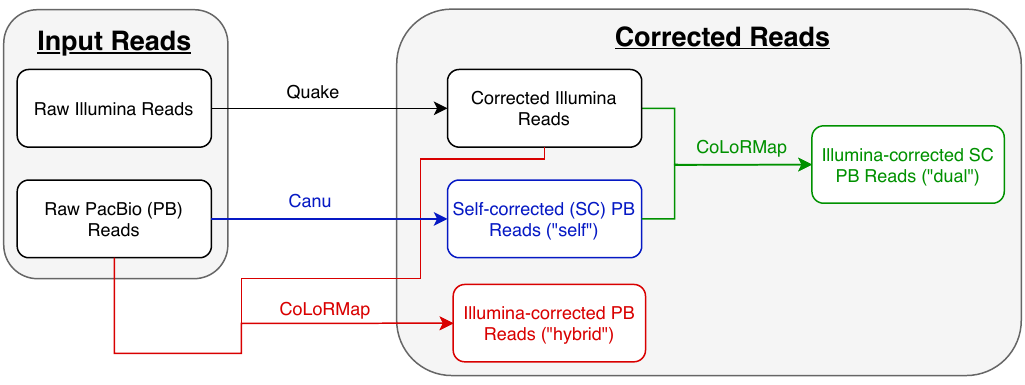

The table and two plots show the NGx and LGx plots where x is a number between 0 and 100 representing the percentage of the genome size. NGx and LGx statistics are similar to the Nx and Lx statistics except they are scaled to the genome size instead of the assembly size. In theory, assemblies improve by maximizing and minimizing the areas under the NG and LG curves, respectively. Plainly, the “dual” correction strategy is superior in terms of continuity.

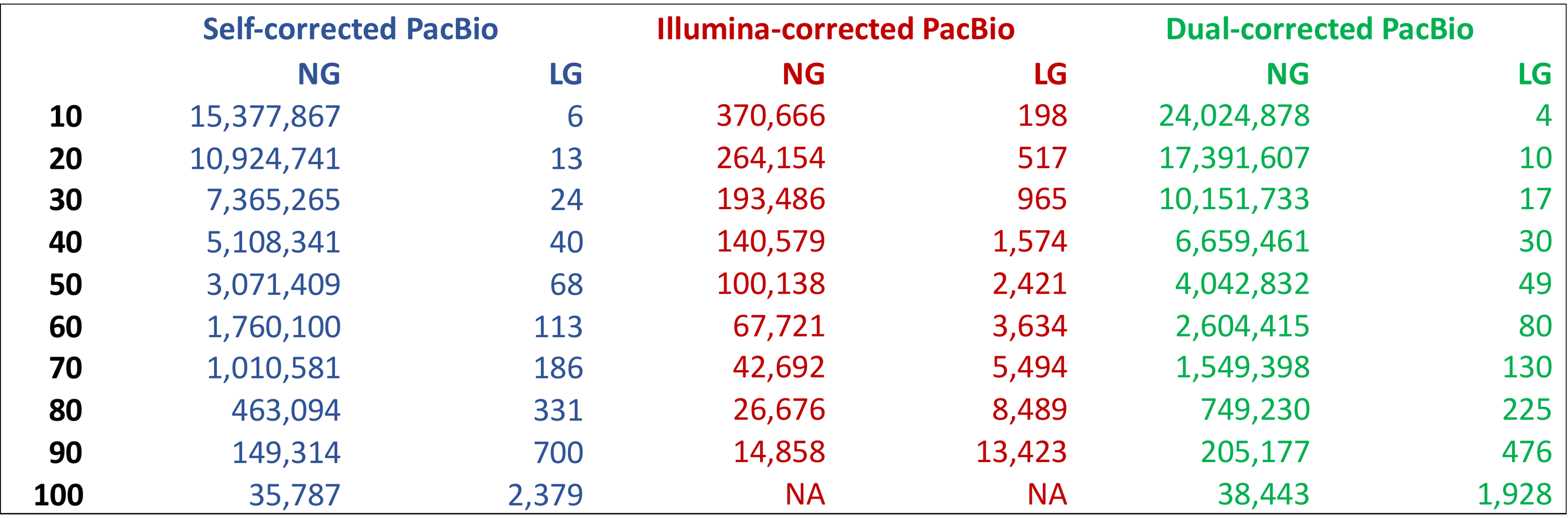

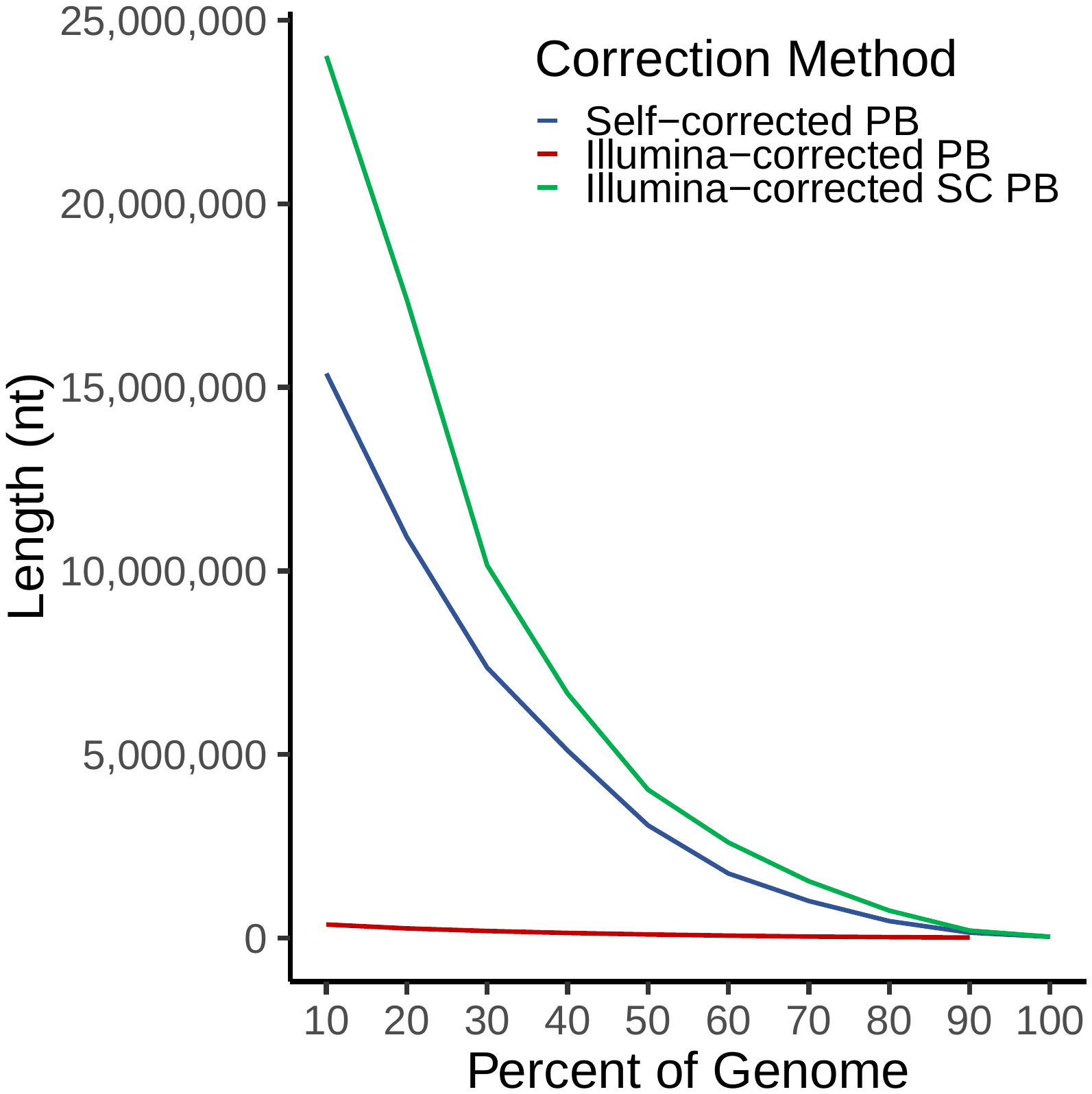

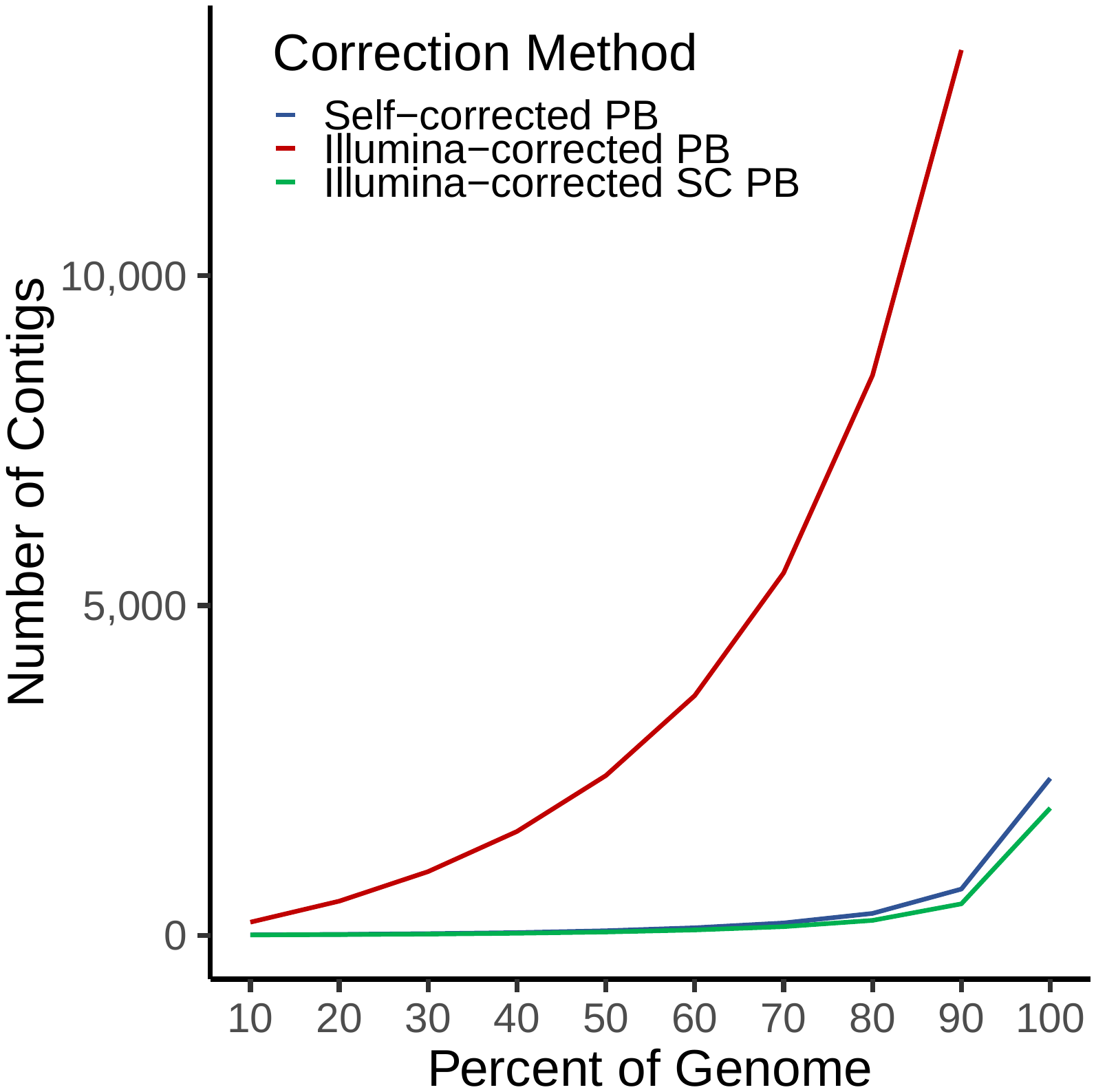

**S.4 – Genome Size Estimation**

ntCard v1.0.1 (Hamid et al. 2017) was used to estimate the k-mer coverage histogram using the following command:

```
ntcard \
    -k 19 \
    -t ${THREADS} \
    -p ${OUTPUT_FILE_BASE_NAME} \
    ${INPUT_FASTQ_FILES[@]}
```

The equation described in the main manuscript was used to determine the genome size from the ntCard output, implemented as a simple AWK program. First, the k-mer coverage histogram must be processed to match the output format of Jellyfish’s histo command (Marcais and Kingsford 2011).

tail -n +3 ${NTCARD_OUTPUT_FILE} \

| tr -d "f" \

> ${HISTO_FILE}

awk -f ${AWK_SCRIPT} ${HISTO_FILE}

Where the AWK program referred to as ${AWK_SCRIPT} is the following:

BEGIN {

x = 0; # x at max y

y = 0; # max y

s = 0; # genome size

}

{

if ($2 >= y) {

y = $2;

x = NR;

}

s += $1 * $2

}

END {

print "peak: " x "," y "; sum: " s "; size: " s / x;

}

**S.5 – Genome Assembly, Polishing, and Scaffolding**

The individual steps of genome assembly, polishing, and scaffolding will each be described separately. Calculation of assembly summary statistics will also be described.

*S.5.1 – Genome Assembly*

The assembly was created with Canu v1.6 (Koren et al. 2017) using the already reads from the “dual” correction strategy using the following command:

```
canu -trim-assemble \
    -s ${SETTINGS_FILE} \
-d ${OUTPUT_DIR_NAME} \
-p ${OUTPUT_PREFIX} \
-pacbio-corrected \
${INPUT_DUAL_CORRECTED_PACBIO_READS_FILE}
```

*S.5.2 – Polishing*

Before polishing the contigs, the corrected Illumina WGS reads required slight modification of the headers because spaces were not allowed. The exact modifications required to make sequence headers match RaCon’s expectations may vary, but the following AWK program worked in our case:

BEGIN {

FS = " ";

}

{

if (NR % 4 == ) {

print $1 "-" substr($2, 1, 1);

} else {

print $0;

}

}

RaCon also required mapping these short reads to the contigs before it would run. The alignments were performed with BWA v0.7.17-r1998 (Li 2013) and converted from SAM format to BAM format using SAMtools v1.6 (Li et al. 2009):

bwa index \

-p ${CONTIGS_INDEX_PREFIX} \

${CONTIGS_FASTQ_FILE}

bwa mem \

-t ${THREADS} \

-p ${CONTIGS_INDEX_PREFIX} \

${ILLUMINA_SHORT_READS_FILE} \

> ${ALIGNMENT_SAM_FILE}

samtools view \

-buS ${ALIGNMENT_SAM_FILE} \

| samtools sort \

-@ ${THREADS} \

> ${ALIGNMENT_BAM_FILE}

Polishing with the corrected Illumina WGS reads using RaCon v1.3.1 (Vaser et al. 2017) was accomplished using the following command:

```
racon \
    --include-unpolished \
--threads ${THREADS} \
${ILLUMINA_SHORT_READS_FILE} \
${ILLUMINA2CONTIGS_ALIGNMENTS_BAM} \
${CONTIGS_FILE} \
> ${POLISHED_CONTIGS_FILE}
```

This process of alignment and polishing was repeated for a second round with the polished output contigs from the first round acting as “unpolished” contigs for the second round.

*S.5.3 – Scaffolding*

The polished contigs were scaffolded in a stepwise fashion using two types of long-range information: Hi-C and RNA-seq reads.

*S.5.3.1 – Hi-C Scaffolding*

The Hi-C data alignments were performed following the Arima Genomics (San Diego, California, USA; https://arimagenomics.com) Mapping Pipeline (https://github.com/ArimaGenomics/mapping_pipeline), which relied on bwa v0.7.17-r1998 (Li 2013), Picard v2.19.2 (Broad Institute 2019), and SAMtools v1.6 (Li et al. 2009). As the pipeline is reasonably well-documented, it will be only summarized here:

1. The assembly (polished contigs) is indexed using SAMtools faidx.
2. The assembly is indexed with bwa index and the Hi-C reads are mapped to the assembly with bwa mem.
3. The alignments are converted from SAM to BAM format with SAMtools view.
4. The 5’ ends are filtered using SAMtools view and the Arima Genomics Perl (https://www.perl.org) script filter_five_end.pl.
5. Paired-end reads are combined into a single file with the Arima Genomics Perl script two_read_bam_combiner.pl and sorted with SAMtools sort. These reads will be treated as single-end hereafter.
6. Read groups are added to the BAM file using Picard AddOrReplaceReadGroups.
7. Merge technical replicates. This step was skipped because no such replicates existed.
8. Duplicates in the BAM file were marked using Picard MarkDuplicates.
9. Merge biological replicates. This step was skipped because no such replicates existed.
10. The final BAM file was indexed with SAMtools index.
11. Stats were reported with the Arima Genomics Perl script get_stats.pl.

Scaffolding was performed on the polished contigs using the final BAM file from the Arima Genomics Mapping Pipeline with SALSA downloaded 29 May 2019 (Ghurye et al. 2017; Ghurye et al. 2019). First, some pre-processing was required with BEDTools v2.28.0 (Quinlan and Hall 2010) to convert the final BAM file from the mapping pipeline to BED format; this was then sorted. The BEDTools, sorting, and SALSA commands are listed here (note that the ${RESTRICTION_ENZYME_SEQ} was GATC):

```
bedtools bamtobed \
    -i ${FINAL_ARIMA_BAM_FILE} \
> ${HIC_BED_FILE}

sort -k 4 \
${HIC_BED_FILE} \
> ${SORTED_HIC_BED_FILE}

run_pipeline.py \
-a ${POLISHED_CONTIGS_FILE} \
-l ${POLISHED_CONTIGS_FAIDX_FILE} \
-b ${SORTED_HIC_BED_FILE} \
-e ${RESTRICTION_ENZYME_SEQ} \
-s ${GENOME_SIZE} \
-m yes \
-o ${OUTPUT_SALSA_DIR}
```

Note that all newly-created gaps from SALSA will all be assigned a length of 500 nucleotides (i.e., 500 Ns in a row). Assuming these are gaps of unknown size, these will ideally be changed to 100 nucleotides for any submissions to GenBank. If you have multiple sources of evidence for gaps (e.g., Hi-C and RNA-seq), you will want to keep track of which gaps were supported by each type of evidence.

*S.5.3.2 – RNA-seq Scaffolding*

The RNA-seq data were aligned using HiSat v0.1.6-beta (Kim et al. 2015), and the alignments were converted from SAM to BAM format and sorted using SAMtools v1.6 (Li et al. 2009). First, the assembly (scaffolds from Hi-C) was indexed with HiSat. For each tissue (i.e., heart, gill, and liver), HiSat aligned reads to the assembly, SAMtools sorted and compressed the output alignments, and Rascaf downloaded June 2018 (Song et al. 2016) computed how scaffolding could be done. The actual scaffolding was done with Rascaf in a single step after all steps had been completed for each tissue. The process is described in the following script:

hisat-build \

${HISAT_IDX_PREFIX} \

${HIC_SCAFFOLDS}

for TISSUE in {gill,heart,liver}

do

RNASEQ_READS_LEFT=${TISSUE}_L.fq.gz

RNASEQ_READS_RIGHT=${TISSUE}_R.fq.gz

ALIGNMENT_SAM=${TISSUE}_aln.sam

hisat \

-p ${THREADS} \

--phred33 -q -t \

-x ${HISAT_IDX_PREFIX} \

-1 ${RNASEQ_READS_LEFT} \

-2 ${RNASEQ_READS_RIGHT} \

-S ${ALIGNMENT_SAM}

samtools view \

-buh ${ALIGNMENT_SAM} \

| samtools sort \

-@ ${THREADS} \

-m ${MEMORY}M \

-O BAM \

-o ${ALIGNMENT_BAM}

rascaf \

-breakN 1 \

-b ${ALIGNMENT_BAM} \

-f ${HIC_SCAFFOLDS} \

-o ${TISSUE}.out

done

rascaf-join \

-r gill.out \

-r heart.out \

-r liver.out \

-o ${OUTPUT_FILE_PREFIX}

Note that the -breakN 1 option breaks all scaffolds at gaps of any size (1 or more Ns) while it determines which sequences it can join. Broken gaps are then restored to their original length and location when additional gaps are added based on the RNA-seq read pairs. If the RNA-seq evidence disagrees with any pre-existing gaps, it will remove them. Also note that newly-created gaps from Rascaf will all be assigned a length of 17 nucleotides (i.e., 17 Ns in a row). For submission to GenBank, these will ideally be changed to 100 nucleotides. If you have multiple sources of evidence (e.g., Hi-C and RNA-seq), you will want to keep track of which gaps were supported by each type of evidence.

*S.5.4 – Assembly Statistics*

Assembly continuity statistics, e.g., N50 and auN (Li 2020), were calculated with caln50 downloaded April 2020 (https://github.com/lh3/calN50) and a custom Python (https://www.python.org) script. caln50 is run using the following simple command:

caln50 \

-s 0.01 \

-L ${GENOME_SIZE} \

${CONTIGS_OR_SCAFFOLDS_FILE} \

> ${STATISTICS_FILE}

The custom Python script is not efficient, but it does calculate Nx, Lx, NGx, and LGx, as well as a few other interesting points about sequences in a fasta file. This script is too long to realistically represent when embedded in the text; it is available on GitHub at https://github.com/pickettbd/basicAsmStatsCalcInPy.

Assembly correctness was assessed using single-copy orthologs with BUSCO v4.0.6 (Simão et al. 2015) and OrthoDB v10 (Kriventseva et al. 2019). The BUSCO config file was the not modified from the default aside from the locations of OrthoDB v10 and the binary executables for BUSCO. It was run based on the following command structure:

busco \

--offline \

--config ${BUSCO_CONFIG_FILE} \

--cpu ${THREADS} \

--in ${CONTIGS_OR_SCAFFOLDS_FASTA} \

--out_path ${OUTPUT_DIR} \

--out ${OUTPUT_FILE_PREFIX} \

--mode genome \

--lineage actinopterygii \

--augustus_species zebrafish

**S.6 – Transcriptome Assembly**

The transcripts were assembled using Trinity v2.6.6 (Grabherr et al. 2011), which depended on Bowtie v2.3.4.3 (Langmead and Salzberg 2012), Jellyfish v2.2.10 (Marcais and Kingsford 2011), salmon v0.12 (Patro et al. 2017), and SAMtools v1.6 (Li et al. 2009):

trinity \

--no_version_check \

--max_memory ${MEMORY} \

--CPU ${THREADS} \

--long_reads ${DUAL_CORRECTED_PACBIO_READS} \

--seqType fq \

--left ${RNASEQ_READS_LEFT} \

--right ${RNASEQ_READS_RIGHT} \

--SS_lib_type FR \

--normalize_max_read_cov 50 \

--normalize_by_read_set \

--min_contig_length 200 \

--output ${TRINITY_OUTPUT_DIR}

Assembly correctness was assessed using single-copy orthologs with BUSCO v4.0.6 (Simão et al. 2015) and OrthoDB v10 (Kriventseva et al. 2019). The command and config file were a match to how BUSCO was run to assess genome assembly correctness, except that the --mode option was transcriptome instead of genome.

**S.7 – Computational Annotation**

The MAKER v3.01.02-beta (Holt and Yandell 2011) pipeline was used to annotate the assembly. With a large enough cluster with MPI support, MAKER runs relatively quickly for each round. The general process was described in prose in the main manuscript, but it can be summarized in outline form here:

I. MAKER round #1
II. *ab initio* gene predictors
    a. AUGUSTUS
    b. GeneMark-ES
    c. SNAP
III. MAKER round #2
IV. *ab initio* gene predictors
    a. AUGUSTUS
    b. SNAP
V. MAKER round #3
VI. MAKER post-processing & functional annotation

As each round of MAKER was run in a nearly identical fashion, the process will be described once, followed by differences between the rounds. Similarly, AUGUSTUS and SNAP will also be described once.

*S.7.1 – MAKER Round #1*

The command to run MAKER is straight-forward, though may vary slightly depending on the implementation of MPI employed by the cluster. The MAKER documentation says to run MAKER with the mpiexec command, but mpirun was successful for our setup. Running MAKER from a working directory on an NFS drive will almost certainly result in failure unless MAKER is directed where to do its work in a non-NFS temporary directory. This required some extra attention to job cleanup on our cluster, but it was successful when we pointed MAKER to the local drives on the nodes on which it was run, which were mounted at /tmp. When calling MAKER from the directory in which the control files exist, the command to start MAKER looks like this:

mpirun maker \

-cpus ${CPUS} \

-TMP ${MAKER_TMP_DIR}

The truly critical parts are in the MAKER control files. Assuming one has a successfully installed and configured version of MAKER available, default control files can be generated in the working directory by running the following command: maker -CTL. No modifications were made to the maker_evm.ctl file. The maker_bopt.ctl file was left unchanged as well. Note that use_rapsearch was set to 0 and blast_type was set to ncbi+. The maker_exe.ctl file was modified as needed only to set correct paths to the executables for MAKER’s dependencies. The following shows the modified or otherwise relevant lines from the maker_opts.ctl file:

```
# genome
genome=/path/to/scaffolds.fa
organism_type=eukaryotic

#re-annotation
maker_gff=
est_pass=0
protein_pass=0
rm_pass=0
model_pass=0
pred_pass=0
other_pass=0

# est/rna-seq
est=/path/to/Trinity/transcripts.fa
est_gff=

# protein homology
protein=/path/to/uniprot_sprot.fa
protein_gff=

# repeat masking
model_org=all
rmlib=/path/to/RepeatModeler/results/assembly-db-families.fa
repeat_protein=/path/to/maker-install-dir/data/te_proteins.fa
rm_gff=
softmask=1

# gene prediction
snaphmm=
gmhmm=
augustus_species=
pred_gff=
model_gff=
run_evm=0
est2genome=1
protein2genome=1
trna=0

# maker behavior
max_dna_len=1000000
min_contig=20000

pred_flank=200
pred_stats=0
AED_threshold=1
min_protein=0
alt_splice=0
always_complete=0
map_forward=0
keep_preds=0

split_hit=10000
min_intron=20
single_exon=0
single_length=250
correct_est_fusion=0
```

Once MAKER has completed, a few MAKER accessory scripts can be run to extract the results from its datastore located at ${PROJECT_DIR}/maker/rnd1/*.datastore. Additional modifications (shown), can also be employed to make output names more palatable. For sake of demonstration, we assume the master datastore index log file is prefixed with scaffolds, and the output base (-o option for fasta_merge) is agloss-rnd1 (*A. glossodonta* round 1)):

cd maker/rnd1/scaffolds.maker.output

fasta_merge \

-o agloss-rnd1 \

-d scaffolds_ master_datastore_index.log

gff3_merge \

-n -s \

-d scaffolds_ master_datastore_index.log \

> agloss-rnd1_noSeq.gff

cd scaffolds_datastore

rename 's/.all.maker./_/' *.fasta # Perl rename, not Linux util

rename 's/fasta/fa/' *.fasta	# Perl rename, not Linux util

awk '{if ($2 == "est2genome") print $0}' \

agloss-rnd1_noSeq.gff \

> agloss-rnd1_est2genome.gff

awk '{if ($2 == "protein2genome") print $0}' \

agloss-rnd1_noSeq.gff \

> agloss-rnd1_protein2genome.gff

awk '{if ($2 ~ "repeat") print $0}' \

agloss-rnd1_noSeq.gff \

> agloss-rnd1_repeats.gff

mv agloss-rnd1*.fa agloss-rnd1*.gff ../..

cd ../../../..

*S.7.2 –* ab initio *Gene Prediction*

Three *ab initio* gene prediction programs were run between MAKER rounds 1 and 2. AUGUSTUS and SNAP can take gene models as input, and they are thus able to be run with new models after rounds 1 and 2 of MAKER in preparation for rounds 2 and 3, respectively. GeneMark-ES does not take gene models as input, and it thus needs to be run only one time.

*S.7.2.1 – GeneMark-ES*

GeneMark-ES required a software key to be run, which can be obtained or re-obtained for free for academic use at any time. GeneMark-ES also requires a configuration file to be run; the default configuration file was used. The following command demonstrates how to run GeneMark-ES:

gmes_petap.pl \

--ES \

--usr_cfg ${COPY_OF_DEFAULT_CONFIG_FILE} \

--cores ${THREADS} \

--sequence ${SCAFFOLDS_ASSEMBLY_FILE}

*S.7.2.2 – AUGUSTUS*

AUGUSTUS training can be handled with BUSCO. Before AUGUSTUS can be trained, configuration files and data from AUGUSTUS and BUSCO will need to be copied to the working directory for this part of the analysis, and the relevant environment variables will need to be reset (which assumes they are properly set in the first place):

```
cp  -r ${AUGUSTUS_CONFIG_PATH} ${PROJECT_DIR}/augustus_config
export AUGUSTUS_CONFIG_PATH=${PROJECT_DIR}/augustus_config

cp  ${BUSCO_CONFIG_FILE} ${PROJECT_DIR}/busco_config.ini
export BUSCO_CONFIG_FILE=${PROJECT_DIR}/busco_config.ini
```

No changes were made to the AUGUSTUS files. The only change made to the BUSCO configuration file was to set download_path=/path/to/odb10 instead of ./busco_download. This is assuming OrthoDB v10 has already been downloaded to that location and that the ‑‑offline flag will be used when running BUSCO. Before training AUGUSTUS, candidate gene regions need to be extracted. This was done with a custom Python script (available at https://github.com/pickettbd/albula-glossodonta_assembly-paper_misc-scripts) and BEDTools v2.28.0 (Quinlan and Hall 2010).

python3 generateBedForMrnaExtraction.py \

maker/rnd1/agloss-rnd1_noSeq.gff \

scaffolds.fa \

candidates-rnd1.bed

bedtools getfasta \

-fi scaffolds.fa \

-bed candidates-rnd1.bed \

-fo candidates-rnd1.fa

AUGUSTUS was trained by running BUSCO with the same command described in the section S.5.4 (i.e., mode=genome, lineage=actinopterygii, augustus_species=zebrafish). To make the AUGUSTUS training parameters generated after running BUSCO available to the next round of MAKER, some post-processing is required:

```
# make dir for final results
mkdir augustus_config/species/agloss

# move to results location
cd "busco-augustus/agloss-rnd1/
run_actinopterygii_odb10/augustus_output/
retraining_parameters/BUSCO_agloss-rnd1"

# rename some files and their references to eachother
rename \ # Perl rename, not Linux util
's/BUSCO_(agloss-rnd1_)/$1/' \
./*

sed \ # gnu sed
-i -r \
's/BUSCO_(agloss-rnd1_)/\1/' \
./agloss-rnd1_parameters.cfg*

# do it again, removing the rnd info
rename \ # Perl rename, not Linux util
's/(agloss)-rnd1)/$1/' \
./*

sed \ # gnu sed
-i -r \
's/(agloss)-rnd1/\1/' \
./*

# copy the files to final results location
cp -f ./* ../../../../../../augustus_config/species/agloss/

# move back to main project dir
cd –
```

*S.7.2.3 – SNAP*

Training with SNAP is much less resource intensive than training AUGUSTUS. Most, if not all, of the commands can reasonably be run “locally” on a login node or other machine. The final output file, genome.hmm, is what will be provided to the next round of MAKER. Inspection of the log files was performed after each step. The process of training SNAP can be described by the following commands:

mkdir -p snap/rnd1

ln -s \

../../maker/rnd1/agloss-rnd1_withSeq.gff \

snap/rnd1/genome.gff

cd snap/rnd1

maker2zff genome.gff

fathom \

genome.ann genome.dna \

-gene-stats \

> gene-stats.log

fathom \

genome.ann genome.dna \

-validate \

> validate.log

fathom \

genome.ann genome.dna \

-categorize 1000 \

> categorize.log

fathom \

uni.ann uni.dna \

-export 1000 -plus \

> export.log

forge \

export.ann export.dna \

> forge.log

hmm-assembler.pl \

genome params \

> genome.hmm

*S.7.3 – MAKER Round #2*

The second round of MAKER was run much the same way as the first, with a few modifications. First, the second round was run in a separate directory: maker/rnd2. The run_evm flag was set to enable MAKER to run EVidenceModeler v1.1.1 (Haas et al. 2008). The control files were copied from the first round and the following changes were made to maker_opts.ctl:

```
# est/rna-seq
est=
est_gff=/path/to/project/maker/rnd1/agloss-rnd1_est2genome.gff

# protein homology
protein=
protein_gff=/path/to/project/maker/rnd1/agloss-rnd1_protein2genome.gff

# repeat masking
model_org=
rmlib=
repeat_protein=
rm_gff=/path/to/project/maker/rnd1/agloss-rnd1_repeats.gff

# gene prediction
snaphmm=/path/to/project/snap/rnd1/genome.hmm
gmhmm=/path/to/project/gmes/output/gmhmm.mod
augustus_species=agloss
run_evm=1
est2genome=0
protein2genome=0
```

Additionally, the same accessory scripts, renaming, etc. was performed after this second round of MAKER as with the first round. The only differences being that rnd1 was replaced with rnd2 in all the commands and names and the awk commands were skipped.

*S.7.4 –* ab initio *Gene Prediction*

Since GeneMark-ES does not take gene models as input, only SNAP and AUGUSTUS could be re-run after MAKER’s second round. Before training them, the models from MAKER were filtered using gFACs v1.1.1 (Caballero and Wegrzyn 2019).

*S.7.4.1 – gFACs Filtering*

In an attempt to improve the quality of gene models being used for this final round of training with AUGUSTUS and SNAP, gFACs was employed to filter out models with single-exon genes, introns shorter than 20bp, etc. The gFACs command and relevant supporting commands (e.g., creating working directories) are shown here:

mkdir -p gfacs/rnd2

ln -s \

../../maker/rnd2/agloss-rnd2_noSeq.gff \

gfacs/rnd2/orig_noSeq.gff

ln -s \

../../assembly/scaffolds.fa \

gfacs/rnd2/assembly.fa

awk \

'BEGIN{x=0;}/^##FASTA/{x=1;}{if(x){print $0;}}' \

maker/rnd2/agloss-rnd2_withSeq.gff \

> gfacs/rnd2/orig_onlySeq.gff

cd gfacs/rnd2

gFACs.pl \

-f "maker_2.31.9_gff" \

-p ./output/agloss-rnd2_noSeq \

--statistics-at-every-step \

--statistics \

--rem-monoexonics \

--min-exon-size 20 \

--min-intron-size 20 \

--min-CDS-size 74 \

--fasta assembly.fa \

--splice-table \

--nt-content \

--canonical-only \

--rem-genes-without-stop-codon \

--allowed-inframe-stop-codons 0 \

--create-gff3 \

--get-fasta-with-introns \

--get-fasta-without-introns \

--get-protein-fasta \

--distributions \

exon_lengths \

intron_lengths \

CDS_lengths \

gene_lengths \

exon_position \

exon_position_data \

intron_position \

intron_position_data \

-O ./output \

orig_noSeq.gff

ln -s \

agloss-rnd2_noSeq_out.gff3 \

output/agloss-rnd2_noSeq.gff

cat \

output/agloss-rnd2_noSeq.gff orig_onlySeq.gff \

> output/agloss-rnd2_withSeq.gff

cd ../..

*S.7.4.2 – AUGUSTUS*

Training AUGUSTUS after the second round of MAKER in preparation for the third round occurred in the same manner as the first time. The exceptions were that (a) the input GFF3 file came from gFACs instead of directly from MAKER, (b) augustus_species=agloss was used instead of augustus_species=zebrafish, and (c) the occurrences of rnd1 in the commands and names were changed to rnd2. The commands are replicated (and appropriately modified) again here:

python3 generateBedForMrnaExtraction.py \

gfacs/rnd2/output/agloss-rnd2_noSeq.gff \

scaffolds.fa \

candidates-rnd2.bed

bedtools getfasta \

-fi scaffolds.fa \

-bed candidates-rnd2.bed \

-fo candidates-rnd2.fa

AUGUSTUS was trained by running BUSCO with the same command described in the section S.5.4 (i.e., mode=genome and lineage=actinopterygii) except that augustus_species=agloss instead of zebrafish. To make the AUGUSTUS training parameters generated after running BUSCO available to the next round of MAKER, some post-processing is required:

```
# move to results location
cd "busco-augustus/agloss-rnd2/
run_actinopterygii_odb10/augustus_output/
retraining_parameters/BUSCO_agloss-rnd2"

# rename some files and their references to each other
rename \ # Perl rename, not Linux util
's/BUSCO_(agloss-rnd2_)/$1/' \
./*

sed \ # gnu sed
-i -r \
's/BUSCO_(agloss-rnd2_)/\1/' \
./agloss-rnd1_parameters.cfg*

# do it again, removing the rnd info
rename \ # Perl rename, not Linux util
's/(agloss)-rnd2)/$1/' \
./*

sed \ # gnu sed
-i -r \
's/(agloss)-rnd2/\1/' \
./*

# copy the files to final results location
cp -f ./* ../../../../../../augustus_config/species/agloss/

# move back to main project dir
cd –
```

*S.7.4.3 – SNAP*

Training SNAP after the second round of MAKER in preparation for the third round occurred in the same manner as the first time. The exceptions were that (a) the input GFF3 file came from gFACs instead of directly from MAKER, (b) the maker2zff command had to be modified, and (c) the occurrences of rnd1 in the commands and names were changed to rnd2. The maker2zff script provided by MAKER that was modified is referred to as maker2zff_v2. The only change required was to use exon instead of CDS on line 142. The commands are replicated (and appropriately modified) again here:

mkdir -p snap/rnd2

ln -s \

../../gfacs/rnd2/output/agloss-rnd2_withSeq.gff \

snap/rnd2/genome.gff

cd snap/rnd2

maker2zff_v2 -n genome.gff

fathom \

genome.ann genome.dna \

-gene-stats \

> gene-stats.log

fathom \

genome.ann genome.dna \

-validate \

> validate.log

fathom \

genome.ann genome.dna \

-categorize 1000 \

> categorize.log

fathom \

uni.ann uni.dna \

-export 1000 -plus \

> export.log

forge \

export.ann export.dna \

> forge.log

hmm-assembler.pl \

genome params \

> genome.hmm

*S.7.5 – MAKER Round #3*

The third round of MAKER was run much the same way as the second, with a few modifications. First, the third round was run in a separate directory: maker/rnd3. The trna flag was used to ensure MAKER ran tRNAscan-SE v1.3.1 (Chan and Lowe 2019). The control files were copied from the second round and the following changes were made to maker_opts.ctl:

```
# gene prediction
snaphmm=/path/to/project/snap/rnd2/genome.hmm
trna=1
```

Additionally, the same accessory scripts, renaming, etc. was performed after this third round of MAKER as with the second round. The only difference being rnd2 replaced with rnd3 in all the commands and names (the awk commands were again skipped).

*S.7.6 – MAKER Post-processing and Functional Annotation*

The structural annotations created by MAKER required some modest post-processing before adding functional annotations. MAKER accessory scripts were used to update sequence names from the long MAKER names to friendlier ones. Other MAKER scripts were used to update the fasta and/or gff3 files with functional annotations found with the BLAST+ Suite v2.9.0 (Altschul et al. 1990; Camacho et al. 2009) and InterProScan v5.45-80.0 (Jones et al. 2014; Mitchell et al. 2019).

```
# create and move to a working dir
mkdir -p maker/post
cd maker/post

# copy the requisite output files
cp ../rnd3/*.gff ../rnd3/*.fa .
cp ../rnd1/agloss-rnd1_{repeats,{est,protein}2genome}.gff .

# remove the rnd info
rename \ # Perl version, not Linux util
's/-rnd[1-3]//' \
*.fa *.gff

# map new ids to MAKER names
NUM_SEQS=`grep -Ev '^#' agloss_noSeq.gff \
| cut -d "\t" -f 9 | tr ';' '\n' \
| cut -d '=' -f 2 | sort -u | wc -l`

maker_map_ids \
--initial=1 \
--prefix=Albula-glossodonta \
--suffix='-?%' \
--iterate=1 \
--justify=${#NUM_SEQS} \
agloss_withSeq.gff \
> identifiers_map.tsv

# rename based on new ids
for FASTA in *.fa
do
cp -f "${FASTA}" "${FASTA%.fa}_renamed.fa"
map_fasta_ids identifiers_map.tsv "${FASTA%.fa}_renamed.fa"
done

for GFF in *.gff
do
cp -f "${GFF}" "${GFF%.gff}_renamed.gff"
map_gff_ids identifiers_map.tsv "${GFF%.gff}_renamed.gff"
done

# prep for functional annotation
cd /path/to/swissprot

makeblastdb \
-dbtype prot \
-in uniprot_sprot.fa \
-input_type fasta \
-title uniprot_sprot \
-hash_index \
-out uniprot_sprot \
-logfile uniprot_sprot_makeblastdb.log

cd –

# do the alignment for func. annot.
blastp \
-task blastp \
-query proteins_renamed.fa \
-db /path/to/swissprot/uniprot_sprot \
-num_threads ${THREADS} \
-max_target_seqs 1 \
-max_hsps 1 \
-evalue 1e-6 \
-outfmt 6 \
-out proteins-x-uniprotSprot_fmt6.tsv

# update the fasta and gff files with func. annots.
for FASTA in *_renamed.fa
do
maker_functional_fasta \
/path/to/swissprot/unitprot_sprot.fa \
proteins-x-uniprotSprot_fmt6.tsv \
${FASTA} \
> ${FASTA%.fa}_putative-function.fa
done

for GFF in *_renamed.gff
do
maker_functional_gff \
/path/to/swissprot/unitprot_sprot.fa \
proteins-x-uniprotSprot_fmt6.tsv \
${GFF} \
> ${GFF%.gff}_putative-function.gff
done

# run interproscan for more func. annots.
interproscan.sh \
-m "standalone" \
-cpu ${THREADS} \
-T "${TMP}" \
-appl "pfam" \
-dp \
-f "TSV" \
-goterms \
-iprlookup \
-pa \
-t "p" \
-i proteins_renamed.fa\
-o proteins-interproscan.tsv

# update the gff files with interproscan results
for GFF in {with,no}Seq_renamed_putative-function.gff
do
ipr_update_gff \
${GFF} \
proteins-interproscan.tsv \
> ${GFF%.gff}_domain-added.gff
done

for GFF in {with,no}Seq_renamed.gff
do
iprscan2gff3 \
proteins-interproscan.tsv \
${GFF} \
> ${GFF%.gff} _visible-iprscan-domains.gff
done

cd ../..
```
