## Additional File 2 for "Genome Assembly of the Roundjaw Bonefish (*Albula glossodonta*), a Vulnerable Circumtropical Sportfish"

— *for* —

**Table S1.** Sampling sites for *A. glossodonta* for population genomic analyses. The number of individuals (N) after data filtering are displayed for each atoll and island group.

| Island Group | Atoll | N (Atoll) | N (Island group) |
| --- | --- | --- | --- |
| Amirantes | St Joseph | 17 | 17 |
| Farquhar | Farquhar | 8 | 17 |
|  | Providence | 9 |  |
| Aldabra | Aldabra | 8 | 14 |
|  | Cosmoledo | 6 |  |
| Mauritius | St. Brandon | 18 | 18 |

**Table S2.** BUSCO statistics for the RNA transcripts and genomic assemblies.

|  | Complete (%) | Complete Single-Copy (%) | Complete Duplicated (%) | Fragmented (%) | Missing (%) | Total |
| --- | --- | --- | --- | --- | --- | --- |
| Transcriptome |  |  |  |  |  |  |
| Trinity Transcripts | 3,144 (86.4) | 1,241 (34.1) | 1,903 (52.3) | 128 (3.5) | 368 (10.1) | 3,640 |
| Genome |  |  |  |  |  |  |
| Canu Contigs | 3,485 (95.7) | 3,081 (84.6) | 404 (11.1) | 22 (0.6) | 133 (3.7) | 3,640 |
| RaCon Polished Contigs | 3,484 (95.7) | 3,076 (84.5) | 408 (11.2) | 22 (0.6) | 134 (3.7) | 3,640 |
| SALSA Scaffolds | 3,480 (95.6) | 3,074 (84.5) | 406 (11.2) | 27 (0.7) | 133 (3.7) | 3,640 |
| SALSA + Rascaf Scaffolds | 3,481 (95.6) | 3,076 (84.5) | 405 (11.1) | 25 (0.7) | 134 (3.7) | 3,640 |

**Table S3.** Input parameters for *ipyrad* used to assemble ddRAD data to the *A. glossodonta* reference genome

| Parameter | Description | Input |
| --- | --- | --- |
| assembly_method | Assembly method | reference |
| datatype | Datatype | ddrad |
| restriction_overhang | Restriction overhang (cut1,) or (cut1, cut2) | TGCAG, CCG |
| max_low_qual_bases | Max low quality base calls (Q<20) in a read | 5 |
| phred_Qscore_offset | phred Q score offset | 33 |
| mindepth_statistical | Min depth for statistical base calling | 6 |
| mindepth_majrule | Min depth for majority-rule base calling | 6 |
| maxdepth | Max cluster depth within samples | 10000 |
| clust_threshold | Clustering threshold for de novo assembly | 0.9 |
| max_barcode_mismatch | Max number of allowable mismatches in barcodes | 0 |
| filter_adapters | Filter for adapters/primers | 2 |
| filter_min_trim_len | Min length of reads after adapter trim | 35 |
| max_alleles_consens | Max alleles per site in consensus sequences | 2 |
| max_Ns_consens | Max N's (uncalled bases) in consensus | 0.05 |
| max_Hs_consens | Max Hs (heterozygotes) in consensus | 0.05 |
| min_samples_locus | Min # samples per locus for output | 10 |
| max_SNPs_locus | Max # SNPs per locus | 0.2 |
| max_Indels_locus | Max # of indels per locus | 8 |
| max_shared_Hs_locus | Max # heterozygous sites per locus | 0.5 |
| trim_reads | Trim raw read edges (R1>, <R1, R2>, <R2) | 0, 0, 0, 0 |
| trim_loci | Trim locus edges (R1>, <R1, R2>, <R2) | 0, 0, 0, 0 |

**Table S4.** Data filtering steps implemented in VCFtools and PLINK after assembly in *ipyrad*.

| SNP Quality Filters |  |
| --- | --- |
| Genotype Calls | Remove individuals missing > 98% genotype calls |
| Indels | Remove indels |
| Read Depth | Remove loci with mean depth > 100 |
| Singletons and minor alleles | Retain sites with a minor allele frequency > 0.05 and minor allele count $\geq$ 2 |
| Biallelic SNPs | Max alleles = 2 |
| Missing Data |  |
|  | Remove loci with genotype call rate < 40% |
|  | Remove individuals missing > 60% genotype calls |
|  | Remove loci with genotype call rate < 60% |
|  | Remove individuals missing > 50% genotype calls |
|  | Remove loci with genotype call rate < 75% |
| Hardy-Weinberg Equilibrium | Remove loci out of HWE (0.05) |
| Linkage Disequilibrium | Remove loci within 1kb windows with r2 > 0.6 |

**Table S5**. Observed heterozygosity (*H*_O_) and expected heterozygosity (*H*_S_) for each island group.

| Island Group | *H*_O_ | *H*_S_ |
| --- | --- | --- |
| Amirantes | 0.2800 | 0.2915 |
| Farquhar | 0.2901 | 0.2946 |
| Aldabra | 0.2589 | 0.2862 |
| Mauritius | 0.2829 | 0.2923 |
